# A negative feedback loop of the TOR signaling moderates growth and enables rapid sensing of stress signals in plants

**DOI:** 10.1101/2020.09.06.284745

**Authors:** Muhammed Jamsheer K, Sunita Jindal, Mohan Sharma, Manvi Sharma, Sreejath Sivaj, Chanchal Thomas Mannully, Ashverya Laxmi

## Abstract

TOR kinase is a central coordinator of nutrient-dependent growth in eukaryotes. Maintaining optimal TOR signaling is critical for the normal development of organisms. However, the mechanisms involved in the maintenance of optimal TOR signaling are currently unknown in plants. In this study, we describe a negative feedback loop of TOR signaling helping in the adaptability of plants in changing environmental conditions. Using an interdisciplinary approach, we identified a plant-specific zinc finger protein FLZ8, as a regulator of TOR signaling in Arabidopsis. In sugar sufficiency, FLZ8 is upregulated by TOR-dependent and –independent histone modifications. FLZ8 negatively regulates TOR signaling by promoting antagonistic SnRK1α1 signaling and bridging the interaction of SnRK1α1 with RAPTOR, a crucial accessory protein of TOR. This negative feedback loop moderates the TOR-growth signaling axis in the favorable condition and helps in the rapid activation of stress signaling in unfavorable conditions establishing its importance in the adaptability of plants.

## INTRODUCTION

Organisms need to coordinate their growth according to nutrient availability. The Target Of Rapamycin (TOR) kinase works as a central regulator of growth in nutrient sufficiency^1,2^. Sugar and other nutrients (nitrogen, amino acids, phosphate, etc.) activate TOR complex 1 (TORC1), which is composed of TOR and the accessory proteins, Regulatory-associated protein of TOR (RAPTOR) and Lethal with Sec Thirteen 8 (LST8)^1^. TORC1 promotes general anabolism and growth^2^. Nutrient starvation especially the sugar starvation, activates AMP-activated protein kinase (AMPK) which reprograms the growth according to the available nutrients^3^. The plant homolog of AMPK is known as Snf1-related protein kinase 1 (SnRK1) is found to be a master regulator of growth in sugar starvation indicating the functional conservation of this pathway^4^. AMPK/SnRK1 phosphorylates RAPTOR leading to the suppression of TOR during starvation^5,6^. Recently, TORC1 was found to suppress AMPK activity through the direct phosphorylation of its α kinase subunit during starvation^7^. Thus, the double-negative feedback loop between TOR and AMPK/SnRK1 coordinates growth in eukaryotes according to nutrient availability^1^.

*TOR* is an essential gene and disruption of *TOR* causes lethal defects in embryo development in plants^8,9^. Further, *TOR* is highly expressed in the primary meristems and drives post-embryonic growth in plants^8^. TOR is also essential for the light and sugar-dependent activation of the meristems through E2F transcription factors^10–12^. Light perception leads to the activation of TOR and photomorphogenesis^11,13^. Thus, TOR works as a critical integrator of light and sugar signals to drive post-embryonic growth. TOR is important for the sugar-dependent hypocotyl elongation in dark indicating its role in skotomorphogenesis^14^.

Hyperactivation of TOR results in severe developmental abnormalities in plants and animals^9,15^. Thus, organisms need to maintain optimal TOR activity for normal development. In animals, TORC1 activates the transcription of *Sestrins* (*SESNs*) which promotes AMPK leading to the attenuation of TORC1 signaling^16,17^. The GATOR1 complex prevents the hyperactivation of TORC1 during amino acid sufficiency^18^. These negative feedback regulators which prevent TORC1 hyperactivity are absent in the plant lineage^19,20^. Thus, plants possibly have evolved plant-specific mechanisms to moderate TOR signaling. However, such regulators are yet to be discovered in plants.

The developing seedlings need to maintain a critical level of TOR activity for cell division and organogenesis. At the same time, they are exposed to various stress factors in the environment. Rapid sensing of stress signals and induction of stress response pathways are crucial for the survival of plants on land^21^. The high TOR activity can attenuate the stress response machinery^1,22^; therefore, plants need to maintain a homeostasis between the activities of TOR and stress signaling to balance growth and stress responses.

In this study, we used an interdisciplinary approach to study the importance of mechanisms that regulate TOR signaling to balance growth and stress signaling in plants. Through a data-driven systems biology approach, we identified potential regulatory factors of TOR signaling. We found that one of such factors identified from the screening, FCS-Like Zinc Finger 8 (FLZ8), acts in a negative feedback loop in TOR signaling and maintains basal SnRK1 and ABA signaling. This regulation was found to be important in moderating growth and rapid activation of stress signaling in plants.

## RESULTS

### Negative feedback regulation of TOR signaling improves fitness of plants in a fluctuating environment

How plants regulate TOR activity to rapidly adjust their growth according to the fluctuations in the environment is not well understood. Negative feedback loops are important in preventing hyperactivation of specific signaling nodes and maintaining homeostasis in cell signaling networks^23,24^. To understand the significance of negative feedback loops in the TOR signaling in plants, we first employed a mathematical modeling approach as it is a powerful tool to decipher the complex and often counter-intuitive dynamics in the cell signaling networks^25^. The mutually antagonistic interaction of TOR and ABA signaling is well understood at the molecular level^22^; therefore, we developed a simple network model for TOR-ABA signaling to understand how plants regulate TOR signaling under different environmental conditions with and without a negative feedback loop denoted as X (Fig. 1A). We used this model to simulate TOR signaling activation and growth progression under energy sufficiency in a system assumed to have baseline TOR and ABA signaling. The negative feedback loop which is activated by TOR under nutrient sufficiency downregulated TOR activity over time leading to the moderation of growth (Fig. 1B, C). We then asked the relevance of such a feedback loop in regulating stress response in changing environmental conditions. In environment, plants that are growing in favorable conditions with high TOR activity can be suddenly exposed to stress conditions. Therefore, we simulated the ABA signaling activation and stress survivability in response to sudden exposure to stress in a system assumed to have high TOR and baseline ABA signaling. The simulation predicted that the presence of X helps in the rapid activation of the stress signaling module which increased the chance of survival (Fig. 1D, E). Taken together, the simulation of the TOR and stress signaling network under different environmental conditions suggest that the negative feedback regulation of TOR signaling would facilitate plants in adjusting their growth in a changing environment.

**Fig. 1:**
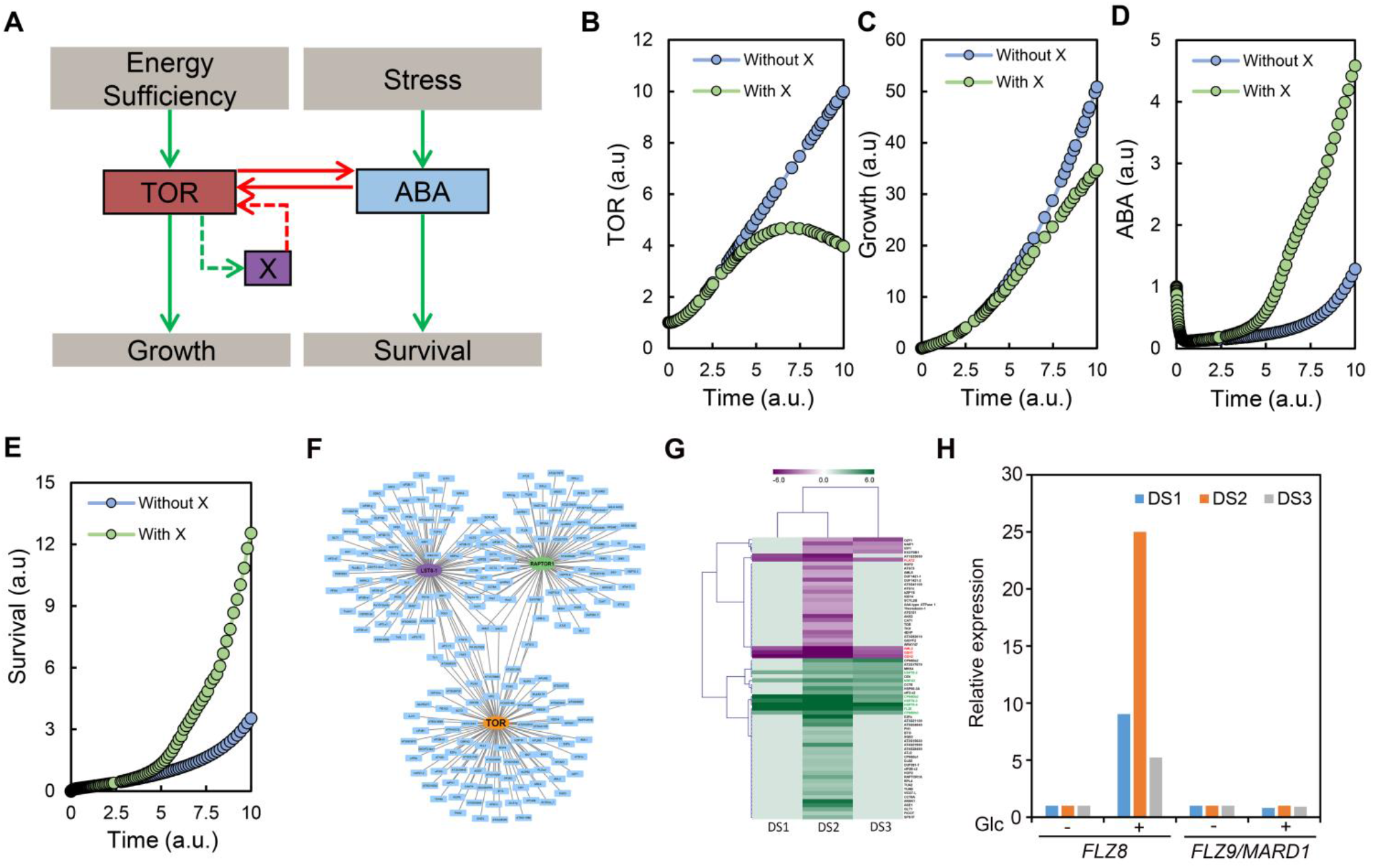
Simulation and data-driven screening for the potential negative feedback regulators of plant TOR signaling. (A) The Network model developed for simulating TOR-ABA signaling with a negative feedback module of TOR (named as X). The green arrows indicate a positive influence and red arrows indicate a negative influence. The arrows connecting X to TOR are represented by dotted arrows. (B) and (C) Simulation of TOR signaling activation and growth under a favorable growth condition with and without X module. (D) and (E) Simulation of ABA signaling activation and survivability in response to sudden exposure to stress with and without X module. (F) The Arabidopsis TOR signaling network which includes identified interacting proteins of TORC components (TOR, RAPTOR1, and LST8-1) and phosphosubstrates of TOR. (G) Heat map of TOR signaling network genes differentially regulated in Glc-transcriptome data sets (DS 1-3). (H) Comparison of transcriptional regulation of *FLZ8* and *FLZ9/MARD1* in response to Glc treatment in the Glc-transcriptome data sets.

### Data-driven screening identifies potential negative feedback regulators of TOR signaling

To identify the negative feedback regulators of TOR, we adopted a data-driven screening approach. We constructed an Arabidopsis TOR signaling network by integrating all the interactors and phosho-substrates of TOR complex identified from individual and high-throughput omics studies (Fig. 1F and Table S1). We screened for the potential negative feedback regulators from this network which comprises 265 genes. To act as a feedback negative regulator of TOR, the gene needs to be activated by TOR. In Arabidopsis, glucose (Glc) treatment rapidly activates TOR^10^. Therefore, to identify the feedback regulators, we checked the transcriptional regulation of all 265 TOR signaling network genes in Glc treatment using three available transcriptome data sets. Among these genes, the expression of 70 genes was found to be differentially regulated upon short-term Glc treatment in one or more data sets (Fig. 1G). 7 genes were found to be induced in all three data sets. Interestingly, one gene of this category named *FLZ8* was found to be a member of the land plant-specific FLZ family implicated in nutrient signaling^26^. FLZ8 physically associates with SnRK1α and β subunits and RAPTOR1B subunit of TOR^27,28^. RAPTOR subunit works as a stop-gate of TORC1 activity^29^. Thus, FLZ8 seems to be a potential factor regulating TOR activity in plants through regulating SnRK1-TOR interaction. A close homolog of FLZ8 named FLZ9/MEDIATOR OF ABA-REGULATED DORMANCY 1 (MARD1) also interacts with SnRK1 subunits and RAPTOR^28,30^ (Fig. 1F and Table S1). However, the expression of *FLZ9/MARD1* was not induced by Glc treatment (Fig. 1H). Further, *FLZ9/MARD1* is a primary ABA and abiotic stress-responsive gene involved in regulating ABA responses^31,32^. Therefore, we focused on *FLZ8* which is a primary Glc-inducible gene for further study. The other 6 genes which were induced in response to Glc treatment in all three transcriptome data sets were found to be heat shock and chaperone proteins and a member of RNA polymerase family (Table S1), thus we did not proceed with them.

### Sugar sufficiency and TOR induce the expression of *FLZ8* by histone modifications

To investigate the role of TOR in regulating the expression of *FLZ8*, we treated seedlings with Torin1, an ATP-competitive inhibitor of TOR^33^. Treatment of Torin1 led to decreased expression of *FLZ8* and two TOR-inducible marker genes *MINICHROMOSOME MAINTENANCE 3* (*MCM3*) and *E2F TARGET GENE 1* (*ETG1*)^10^ (Fig. 2A). Further, the expression of *FLZ8* and marker genes were found to be increased in two TOR overexpression lines (TOR OE1/GK-166C06 and TOR OE2/GK-548G07)^34^ (Fig. 2B). These results collectively suggest a positive correlation between TOR signaling and the expression of *FLZ8*.

**Fig. 2:**
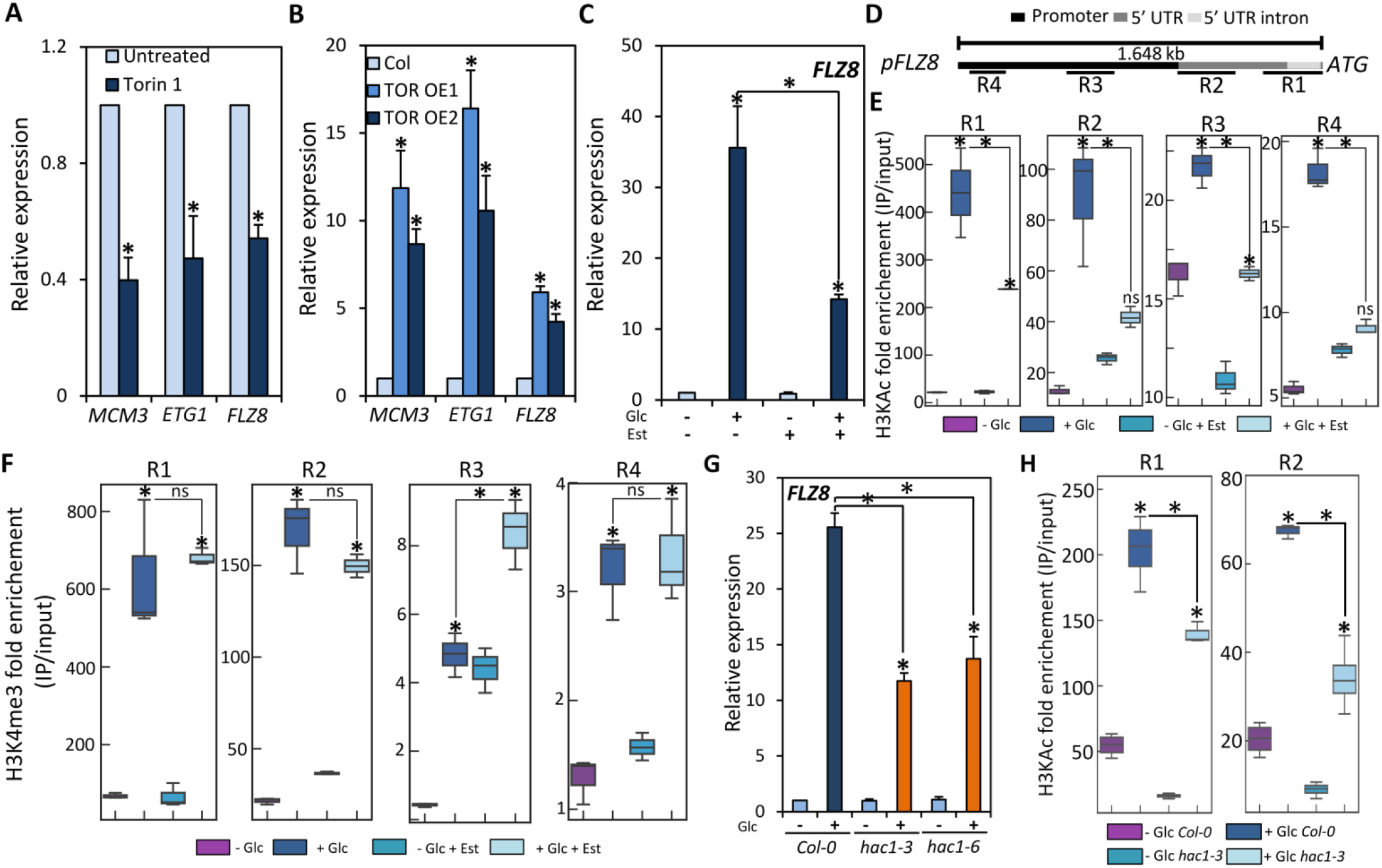
TOR-dependent and -independent pathways direct specific histone modifications to promote the expression of *FLZ8* during sugar sufficiency. (A) Expression of *MCM3*, *ETG1*, and *FLZ8* in response to Torin 1 (10 μM) treatment (one-way ANOVA, *p ≤ 0.05, Bonferroni post-hoc test). (B) Expression of *MCM3*, *ETG1*, and *FLZ8* in TOR overexpression (*TOR OE*) lines (one-way ANOVA, *p ≤ 0.05, Bonferroni post-hoc test). (C) Expression of *FLZ8* in *tor-es1* line (growing in 0 or 10 μM estradiol) treated with 0 or 170 mM Glc (two-way ANOVA, *p ≤ 0.05, Bonferroni post-hoc test). (D) Promoter and upstream regulatory regions of *FLZ8.* The regions (R1 to R4) tested in ChIP-qPCR are indicated. (E) Histone acetylation (H3KAc) status in the upstream region of *FLZ8* in *tor-es1* line (growing in 0 or 10 μM estradiol) treated with 0 or 170 mM Glc (two-way ANOVA, *p ≤ 0.05, Bonferroni post-hoc test). (F) Histone methylation (H3K4me3) status in the upstream region of *FLZ8* in *tor-es1* line (growing in 0 or 10 μM estradiol) treated with 0 or 170 mM Glc (two-way ANOVA, *p ≤ 0.05, Bonferroni post-hoc test). (G) Expression of *FLZ8* in *hac1* mutant lines treated with 0 or 170 mM Glc (n=3) (two-way ANOVA, *p ≤ 0.05, Bonferroni post-hoc test). (H) Histone acetylation (H3KAc) status in the upstream region of *FLZ8* promoter in *hac1-3* treated with 0 or 170 mM Glc (two-way ANOVA, *p ≤ 0.05, Bonferroni post-hoc test). The error bars of bar graphs indicate SE.

To investigate the role of TOR in Glc-dependent induction of *FLZ8*, we used estradiol (Est)-inducible *TOR* RNAi line (*tor-es1*)^35^. The short-term (3 h) Glc treatment led to a strong induction of *FLZ8* in *tor-es1* which was significantly reduced upon the Est treatment (Fig. 2C). It is known that TOR associates with epigenetic factors and regulates histone modifications to control gene expression in response to sugar sufficiency^36^. Histone 3 lysine acetylation (H3KAc) and Histone 3 lysine-4 tri-methylation (H3K4me3) are generally associated with the activation of gene expression^37^. To identify the possible role of TOR and histone modification in regulating the rapid transcriptional upregulation of *FLZ8* in response to Glc treatment, we performed a ChIP-qPCR using anti-H3KAc and anti-H3K4me3 antibodies in *tor-es1* treated with Glc or Glc and Est. Glc treatment led to enhanced enrichment of H3KAc and H3K4me3 in the upstream regulatory region of *FLZ8* with more enrichment in the region closer to the transcriptional start site (TSS) (Fig. 2D-F). TOR inhibition through Est treatment reduced the Glc-dependent recruitment of H3KAc (Fig. 2E). Conversely, the recruitment of H3K4me3 was not reduced in response to TOR inhibition (Fig. 2F). Taken together, the qPCR and ChIP-qPCR results indicate that TOR promotes the recruitment of H3KAc in the promoter region of *FLZ8* leading to the activation of its transcription under Glc sufficiency. TOR phosphorylates and activates a CREB-binding protein/p300-type Histone Acetyltransferase in mammals^38^. Its Arabidopsis orthologue *Histone Acetyltransferase1 (HAC1*) acetylates genes in response to sugar treatment to induce their expression^39^. We found that the induction of *FLZ8* level in response to Glc is reduced in *hac1* mutants indicating the possible role of HAC1 in regulating the acetylation of *FLZ8* promoter in Glc sufficiency (Fig. 2G). To test this hypothesis, we performed a ChIP-qPCR using the anti-H3KAc antibody in the *hac1* mutants. We found that the Glc-dependent induction of H3KAc in the *FLZ8* promoter is significantly reduced in the *hac1* mutants (Fig. 2H; Fig. S1). Collectively, our results indicate that the TOR-dependent H3KAc and TOR-independent H3K4me3 in the upstream regulatory regions of *FLZ8* induce its expression during sugar and energy sufficiency.

TOR phosphorylates E2Fa, which in turn binds to the promoters of target genes to drive their expression^10,40^. To identify the possible role of E2Fa in regulating the expression of *FLZ8*, we analyzed the presence of the E2Fa binding sites in the promoter region of *FLZ8*. We didn’t find any typical E2Fa binding sites described previously^41^ in the *FLZ8* promoter. However, we found sites with similarity to the typical E2Fa binding sites. Therefore, we performed a ChIP-qPCR using the anti-E2Fa antibody to check the binding of E2Fa on the promoter of *FLZ8*. We used the *e2fa* mutant as a negative control along with WT and enrichment was analyzed in different promoter regions of FLZ8 and a previously identified E2Fa target *MCM5*^42^ (Fig. S2A). In line with the previous report, specific enrichment of E2Fa was observed in the promoter of *MCM5* in WT which was significantly reduced in *e2fa* (Fig. S2B). However, no enrichment was observed in any of the four regions in the 1.6 kb upstream region of *FLZ8* in WT with similar signal intensities in both WT and *e2fa* (Fig. S2C). These results indicate that E2Fa doesn’t bind to the promoter of *FLZ8* to drive its expression in Glc sufficiency. Further, we found that the expression of *FLZ8* was not significantly altered in the *e2fa* (Fig. S2d). Taken together, these results clearly indicate that the E2Fa pathway is not involved in the TOR-mediated induction of *FLZ8* level during sugar sufficiency.

Along with light and sugar, mineral nutrient sufficiency (nitrogen, phosphate, sulfur, etc.) also activates TOR^10,13,43,44^. As *FLZ8* levels are rapidly induced in response to short-term Glc treatment in a TOR-dependent manner (Fig. 2), we aimed to identify whether its expression is also induced in response to mineral nutrient (nitrogen, phosphate, iron, sulfur, and potassium) treatment using publically available transcriptome data. We found a clear positive correlation between *FLZ8* transcript level and sugar sufficiency (Fig. S3A). However, such correlation was absent between induction of *FLZ8* transcript level and the other nutrients. Further, the expression of *FLZ8* was found to be upregulated in response to high light treatment and illumination of etiolated plants (Fig. S3B). The expression of *FLZ8* is downregulated under sugar starvation which was more pronounced in the overexpression of *SnRK1α1*, the major kinase of SnRK1 (Fig. S4A). Further, *SnRK1α1* overexpression abolished the induction of *FLZ8* expression in response to sugar treatment (Fig. S4B). Taken together, our results indicate that the Glc-TOR signaling is the specific upstream activator of *FLZ8.* SnRK1 signaling which operates majorly during sugar starvation suppresses *FLZ8* indicating that *FLZ8* level is strongly correlated with the changes in the TOR and SnRK1 signaling in plants.

### TOR signaling is negatively regulated by FLZ8 in plants

Considering the involvement of *FLZ8* in TOR signaling network, we sought to investigate its role in growth and development. We identified two mutants of *FLZ8* (Fig. 3A, B). We analyzed the phenotype of these *flz8* mutants from the early seedling stage as TOR is an important regulator of seedling development^45^. Cotyledon opening is a crucial TOR-mediated photomorphogenic response during the early seedling development^13^. Light reception promotes auxin signaling which activates the TOR pathway leading to the promotion of translation and cotyledon opening. Defects in TOR and its downstream signaling partners lead to the suppression of cotyledon opening^13^. Under favorable light (60 μmol m^−2^ sec^−1^), *flz8* mutant lines showed enhanced cotyledon opening (Fig. 3C). Further, *flz8* seedlings showed enhanced growth with significantly longer primary roots and increased lateral root number, and biomass (Fig. 3D, E). In response to sugar sufficiency, TOR activates root meristem to drive postembryonic root development. In line with this, TOR inactivation leads to a reduction in the cell cycle activity and root meristem size^42^. We found that *flz8* alleles show a clear increase in the length and diameter of the root meristem (Fig. 3F). Further, *in situ* monitoring of cell cycle activity using 5-ethynyl-2′-deoxyuridine (EdU; an analog of thymidine) staining identified meristem activity in a larger area in the roots of *flz8* lines (Fig. 3G). Therefore, *flz8* lines displayed enhanced root growth per day in the growth kinetics analysis (Fig. 3H). Similar to the seedling stages, mutants consistently accumulated more biomass in different stages of rosette development (Fig. 3I). Collectively, the phenotypic analysis identified that the loss of *FLZ8* causes meristem hyperactivation leading to enhanced growth and biomass.

**Fig. 3:**
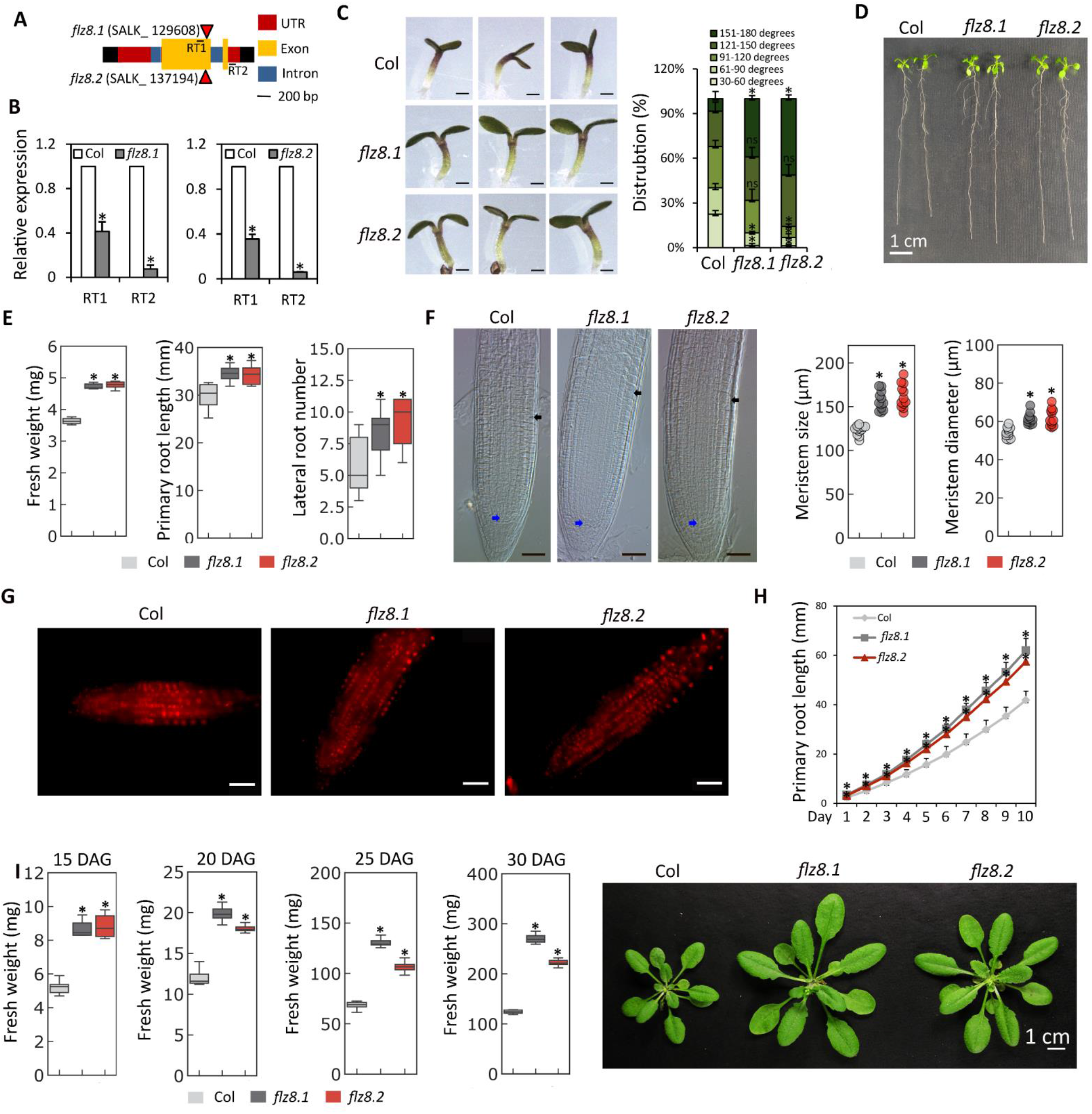
Meristem activity and growth are enhanced in *flz8* mutants. (A) Gene structure of *FLZ8* showing the T-DNA insertion site in *flz8.1* and *flz8.2* alleles. The position of primer sets (RT1 and RT2) used for qPCR is also shown in the gene structure. (B) Expression of *FLZ8* in mutant lines (one-way ANOVA, *p ≤ 0.05, Bonferroni post-hoc test). (C) Cotyledon angle of WT and *flz8* lines at 3 DAG. (D) and (E) Phenotype and measurement of biomass, primary root length, and lateral root number of WT and *flz8* lines at 10 DAG. (F) Root meristem size (length and diameter) of WT and *flz8* lines at 5 DAG. Blue arrow points to the quiescent centre and black arrow points to the junction between meristem and elongation zone (G) Edu (5-ethynyl-2′-deoxyuridine) staining showing root meristem activity of WT and *flz8* lines at 5 DAG. (H) Root growth kinetics in WT and *flz8* lines. (I) Measurement of rosette biomass at different rosette stages and phenotype of WT and *flz8* lines at 30 DAG (one-way ANOVA, *p ≤ 0.05, Bonferroni post-hoc test).

The phenotypic alterations identified in the *flz8* mutants are typically associated with TOR hyperactivation^10–12^. The ribosomal S6 Kinases (S6Ks) are conserved phosho-substrates of TOR and their phosphorylation status is widely used as a readout of TOR activity^11,26,35,43^. Immunoblotting using specific antibodies identified a significant increase in the phosphorylated form of S6Ks in both *flz8* lines indicating the enhancement of TOR activity (Fig. 4A). However, the transcript level of TOR complex components and the two paralogs of *S6Ks* were found to be unaltered indicating that FLZ8 negatively regulates the TOR signaling through regulating its activity (Fig. S5). Changes in the TOR activity can lead to alteration in the sensitivity towards TOR inhibitors^46,47^. We found that *flz8* alleles showed more resistance to TOR inhibitor AZD-8055 at moderate concentrations (0.25 and 0.5 μM) (Fig. 4B). TOR promotes sugar-induced hypocotyl elongation in dark^14^. In line with their high TOR activity, *flz8* alleles showed enhanced sucrose-induced elongation of hypocotyls in dark (Fig. 4C). Collectively, these results indicate that loss of FLZ8 leads to TOR hyperactivation leading to enhanced growth in favorable growth conditions.

**Fig. 4:**
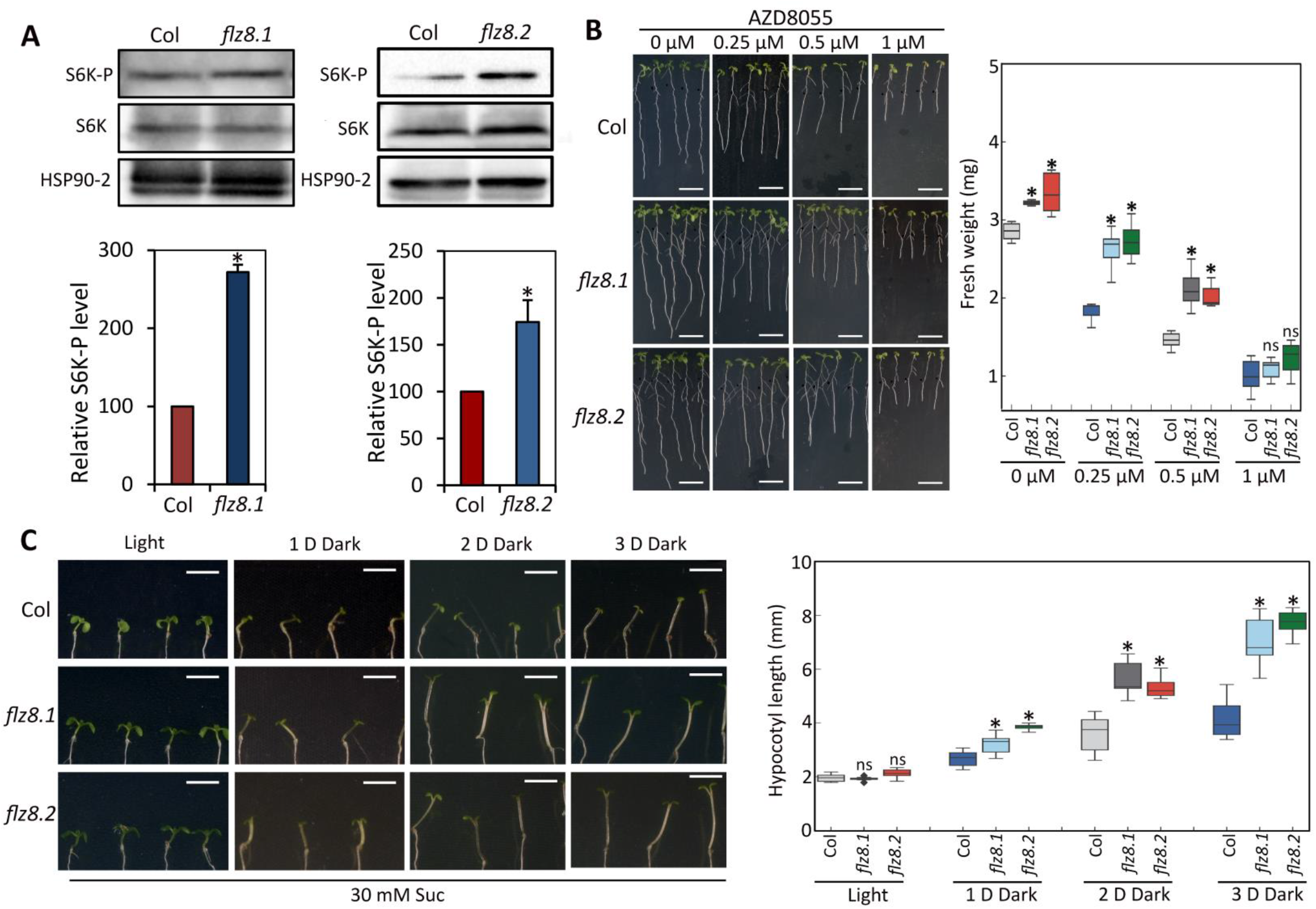
TOR signaling is hyperactivated in *flz8* mutants. (A) Level of total and phosphorylated S6Ks (Thr-449 in S6K1 and Thr-455 in S6K2) in WT and *flz8* mutants at 10 DAG. Relative level of S6K-P was quantified in comparison to total S6K (one-way ANOVA, *p ≤ 0.05, Bonferroni post-hoc test). (B) Phenotype and fresh weight of WT and mutant lines in different concentrations of TOR inhibitor AZD8055 (two-way ANOVA, *p ≤ 0.05, Bonferroni post-hoc test). (C) Sucrose-dependent hypocotyl elongation in WT and *flz8.1* in response to dark treatment for 1, 2 and 3 days (two-way ANOVA, *p ≤ 0.05, Bonferroni post-hoc test).

*TOR* is highly expressed at the early developmental stages and plays an important role in early seedling development^8,10–12^. We used the *pFLZ8:: GUSA* lines to monitor the overlap of *FLZ8* expression domain with *TOR*. Interestingly, these lines showed maximum GUS activity at early seedling stages (2 and 3 DAG). The promoter activity progressively decreased in the later stages (Fig. S6A). In adult stages, GUS activity was low in young leaves and it progressively increased in older and senescing leaves which are characteristic of the *FLZ* family (Fig. S6B)^26,48^. GUS activity was also observed specifically in the developing pollen (Fig. S6C, D). These results suggest that similar to *TOR*, *FLZ8* is also highly expressed during the early seedling stages.

### FLZ8 associates with critical domains of SnRK1α1 and RAPTOR1B and promotes the SnRK1α1-RAPTOR1B interaction

We sought to investigate the mechanism of FLZ8-mediated negative regulation of TOR. FLZ8 physically associates with SnRK1, the prominent negative regulator of TOR signaling. Further, based on the promiscuous interaction of FLZ proteins with SnRK1 and shared interacting proteins, it is hypothesized that FLZ proteins might be working as scaffold proteins that help in the recruitment of phospho-substrates to the SnRK1 complex^27,49^. To understand the possible role of FLZ8 in SnRK1 signaling, we used the yeast-two-hybrid (Y2H) assay to map the interaction site of FLZ8 in SnRK1α1 (Fig. 5A). Mapping of the interaction site identified that FLZ8 interacts strongly with both the catalytic domain (CD) and the regulatory domain (RD). The RD region harbors Ubiquitin-associated domain (UBA) and the αC-terminal domain (αCTD) which is important for the heterotrimeric enzyme complex formation^1^. When the interaction site in the RD region was mapped, FLZ8 specifically interacted with the UBA domain (Fig. 5A). The UBA domain was recently found to be important for the catalytic activity of SnRK1α1^50^. In a large-scale interactome study, FLZ8 was found to be interacting with RAPTOR1B^28^. To confirm this interaction, we used Y2H assay with an appropriate control experiment which identified the strong interaction of FLZ8 with RAPTOR1B (Fig. 5B). Further, we mapped the interaction site of FLZ8 in RAPTOR1B in which FLZ8 showed specific interaction with Raptor N-terminal CASPase-like (RNC) domain and WD40 repeats located in the C-terminus (Fig. 5C). In mammals, these domains are very critical for TORC1 activity as mutagenesis in these domains abolish the RAPTOR-TOR association^51^. Thus, our mapping analysis identified that FLZ8 makes extensive contacts with SnRK1α1 and RAPTOR1B at multiple sites which are critical for the SnRK1 and TORC1 activity respectively.

**Fig. 5:**
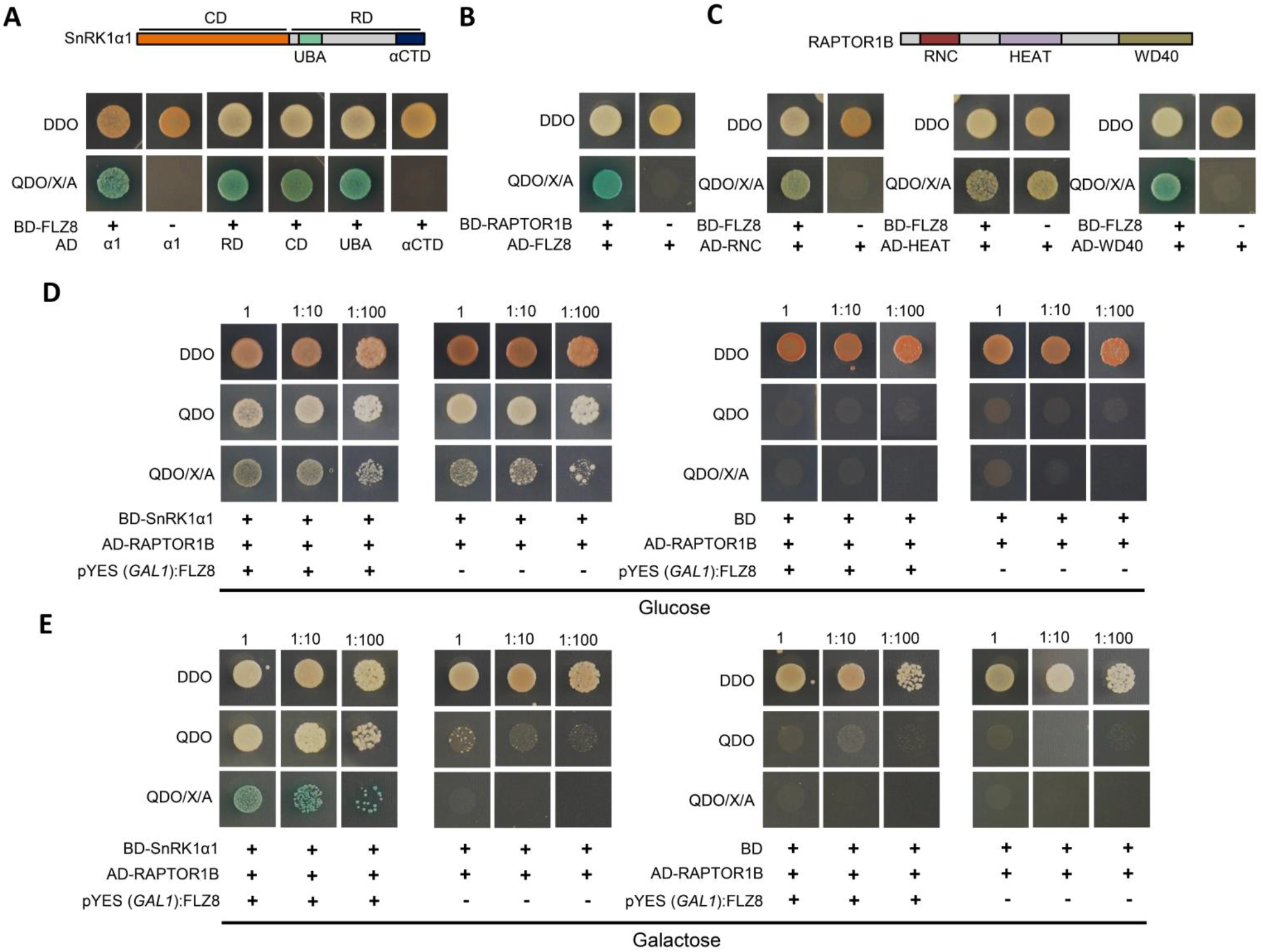
FLZ8 promotes the interaction of SnRK1α1 with RAPTOR1B. (A) Domain organization of SnRK1α1 and mapping of FLZ8-interaction sites in SnRK1α1 by Y2H analysis. (B) Interaction of FLZ8 with RAPTOR1B. (C) Domain organization of RAPTOR1B and mapping of FLZ8-interaction sites in RAPTOR1B. (D) and (E) Analysis of the interaction of SnRK1α1 and RAPTOR1B in the presence of FLZ8 by Y3H analysis in glucose and galactose. For Y2H, the cotransformed yeast cells carrying different AD and BD construct combinations were spotted on synthetic double dropout (DDO; -Trp/-Leu) and quadruple dropout medium supplemented with 40 μg/mL X-α-Gal and 200 ng/mL AbA (QDO/X/A; -Ade/-His/-Leu/-Trp/+ X-α-Gal/+ AbA). For Y3H, the cotransformed yeast cells carrying different AD and BD construct combinations along with pYES (*GAL1*) FLZ8) or pYES (*GAL1*) growing in glucose or galactose were spotted on DDO medium and weak (QDO) and strong (QDO/X/A) interaction screening medium.

The mapping results indicate the potential role of FLZ8 as a scaffold protein which mediates the interaction of SnRK1α1 with RAPTOR1B. To test this hypothesis, we employed a yeast-three-hybrid (Y3H) assay in which along with SnRK1α1 with RAPTOR1B, FLZ8 was expressed under a glucose-repressed and galactose-inducible promoter (*GAL1*). The expression of FLZ8 in galactose strongly enhanced the interaction of SnRK1α1 with RAPTOR1B which was found to be absent in the cells growing in glucose (Fig 5D, E). Thus, these results identified the molecular function of FLZ8 as a scaffold protein that connects TOR and SnRK1 complexes in plants.

### SnRK1 signaling is positively regulated by FLZ8 in plants

In protein-protein interaction assays, we found that FLZ8 interacts with the UBA domain of SnRK1α1 which is generally involved in preventing protein ubiquitination^52^. Thus, we hypothesized that FLZ8 might be involved in the regulation of SnRK1α1 stability. To test this possibility, we estimated the endogenous level of SnRK1α1 in *flz8* mutants at the seedling stage growing under the normal growth conditions using a specific antibody. Immunoblot analysis identified a clear reduction in the level of endogenous SnRK1α1 in the mutants (Fig. 6A). The transcript level of *SnRK1α1* and its paralog *SnRK1α2* were unaltered in the *flz8* mutants (Fig. S7). Thus, FLZ8 promotes SnRK1 signaling through maintaining the level of SnRK1α1.

**Fig. 6:**
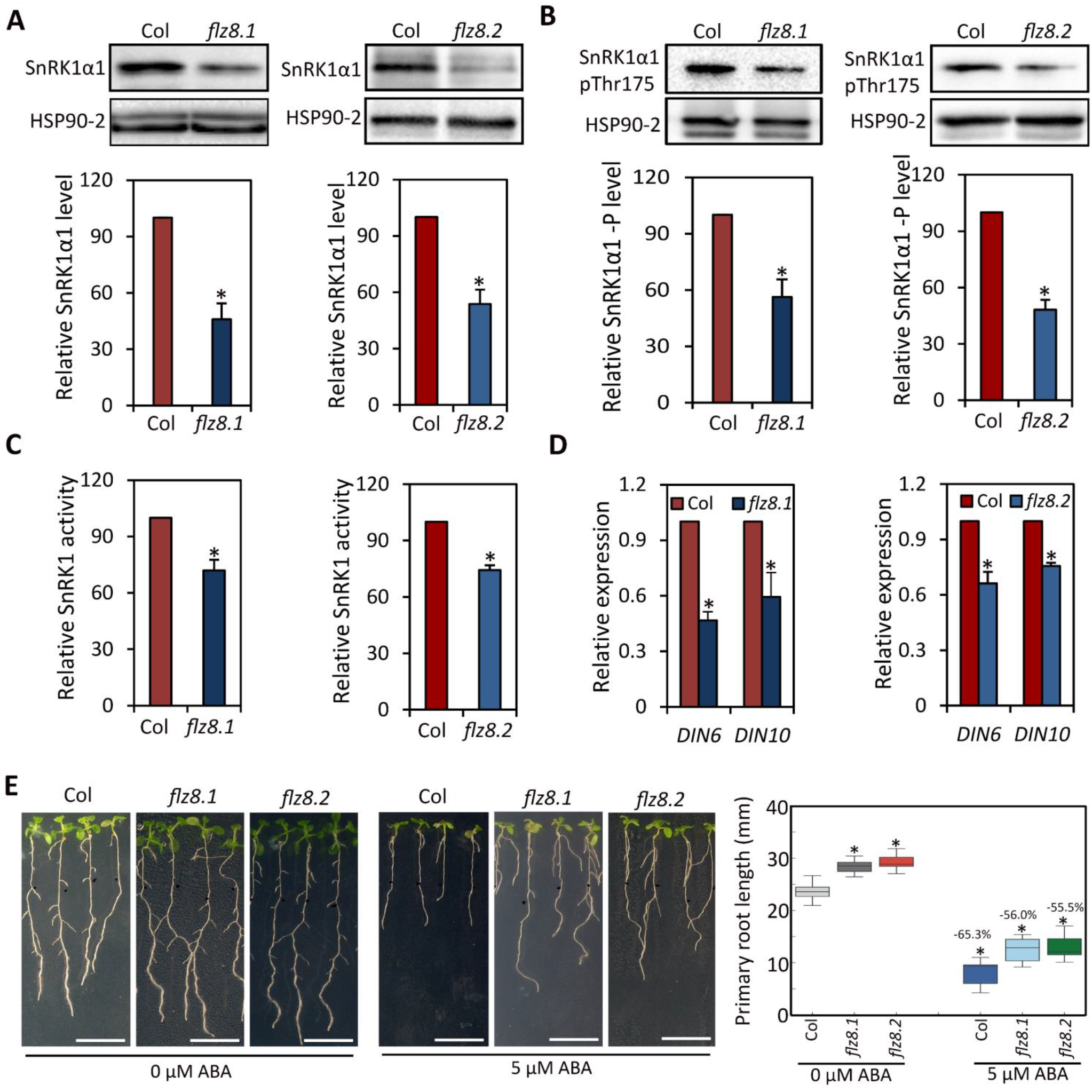
SnRK1 signaling is attenuated in *flz8* mutants. (A) Level of SnRK1α1 in WT and *flz8* mutants at 10 DAG. Relative level of SnRK1α1 was quantified in comparison to HSP90-2 level (one-way ANOVA, *p ≤ 0.05, Bonferroni post-hoc test). (B) Level of phosphorylated SnRK1α1 (Thr-175) in WT and *flz8* mutants at 10 DAG. Relative level of SnRK1α1-P was quantified in comparison to HSP90-2 level (one-way ANOVA, *p ≤ 0.05, Bonferroni post-hoc test). (C) SnRK1 activity in 10 DAG *flz8* mutants in comparison to WT in AMARA peptide assay (one-way ANOVA, *p ≤ 0.05, Bonferroni post-hoc test). (D) Expression of SnRK1-activity reporter genes (*DIN6* and *DIN10*) in 10 DAG *flz8* mutants in comparison to WT (one-way ANOVA, *p ≤ 0.05, Bonferroni post-hoc test). (E) Phenotype and primary root length of WT and *flz8* mutants in response to ABA treatment for 5 days (two-way ANOVA, *p ≤ 0.05, Bonferroni post-hoc test).

Phosphorylation at the Thr-175 in T-loop is essential for the activity of SnRK1^4^. Therefore, we tested whether the decrease in the total pool of SnRK1α1 in the *flz8* mutants results in a reduction in the active pool of kinase using an antibody that detects Thr-175 phosphorylated form of SnRK1α1. In the immunoblotting, *flz8* seedlings growing under normal growth conditions showed a clear reduction in the level of Thr-175 phosphorylated SnRK1α1 (Fig. 6B). We further analyzed its effect on endogenous SnRK1 activity using AMARA peptide assay. SnRK1 activity was found to be significantly decreased in the *flz8* seedlings in comparison to the WT (Fig. 6C). SnRK1 promotes the expression of *Dark Induced 6* (*DIN6*) by phosphorylating bZIP63^53^, and *Dark Induced 10* (*DIN10*)^4^. The expression of *DIN6* and *DIN10* is positively correlated with SnRK1 activity and thus widely used as SnRK1 signaling readout^4,53–55^. In line with the reduced SnRK1 activity in the *flz8* mutants, we found a significant downregulation of *DIN6* and *DIN10* transcript levels in the mutants (Fig. 6D). Hyperactivation of SnRK1 signaling leads to ABA hypersensitivity including enhanced inhibition in the primary root growth in response to ABA treatment^26,56^. Therefore, we tested the ABA sensitivity of *flz8* mutants at the seedling stage and found that ABA treatment led to reduced inhibition of primary root growth in the mutants (Fig. 6E). Autophagy is positively mediated by SnRK1 and negatively regulated by TOR signaling^57–59^. Using a specific antibody, we estimated the endogenous autophagy level of *flz8* mutants through assaying the level of ATG8a lipidation with phosphatidylethanolamine (ATG8a-PE), which is important for the autophagosome formation^60^. The immunoblotting analysis revealed a clear reduction in the level of ATG8a-PE in the *flz8.1* line (Fig. S8). Taken together, our results establish FLZ8 as a positive regulator of SnRK1 signaling in normal growth conditions by maintaining the level of SnRK1α1. Loss of FLZ8 leads to reduced endogenous SnRK1α1 level which culminates in reduced SnRK1 activity leading to changes in gene expression, ABA response, and autophagy.

### The FLZ8-mediated negative feedback module of TOR works as a moderator of growth during sugar sufficiency

Our findings show that the level of FLZ8 is a critical factor balancing the TOR-SnRK1 signaling in plants. Induction of FLZ8 during sugar sufficiency by TOR-dependent and independent pathways leads to the maintenance of SnRK1 signaling via stabilization of SnRK1α1 and the promotion of SnRK1α1-RAPTOR1B association. During sugar starvation, SnRK1 signaling downregulates the FLZ8 level indicating that this module mainly operates in TOR-SnRK1 signaling during sugar sufficiency conditions. To better understand the importance of this negative feedback module on TOR-SnRK1 signaling and growth, we developed a dynamic network model according to our results (Fig. S9A). Using this model, we simulated the effect of FLZ8-mediated regulation on TOR, SnRK1, and biomass accumulation in favorable growth conditions (i.e. energy sufficiency). Simulation of the network under energy sufficiency predicted that FLZ8-mediated regulation moderates TOR signaling and growth by keeping a basal level of SnRK1 signaling (Fig. S9B-D). These results are consistent with our experimental results in which we found hyperactivation of TOR signaling and reduction in the basal SnRK1 signaling in the *flz8* mutants growing in optimal light and sugar (30 mM Suc) level (Fig. 4 and 6). To further clarify the role of FLZ8 in sugar availability-dependent growth, we transferred the 5DAG WT, *flz8.1*, *tori* (*tor 35-7*) seedlings to sugar starvation (8 mM Suc) and sugar abundant (60 mM Suc) growth regimes and the effect was assessed after 5 days. The growth and biomass of all lines were comparable to WT at 8 mM Suc indicating the major role of TOR and FLZ8 in regulating growth under sugar sufficiency (Fig. S9E and F). However, significant differences in growth and biomass were observed at 60 mM Suc. Consistent with the role of TOR signaling in plant growth and biomass accumulation in nutrient-rich conditions, the *tori* line showed reduced overall growth and biomass accumulation compared to WT. In comparison, *flz8.1* showed more vigorous growth with more biomass in 60 mM Suc (Fig. S9E and F). Taken together, the simulation and experimental results establish FLZ8 as a critical regulator of sugar-dependent growth through modulating the TOR-SnRK1 signaling.

### The FLZ8-mediated negative feedback module of TOR helps in the rapid induction of SnRK1 and ABA signaling during stress

Our results indicate that loss of FLZ8 leads to the downregulation of basal SnRK1 signaling which culminates in TOR hyperactivation and negative regulation of autophagy. SnRK1 signaling and autophagy are intimately linked with stress signaling in plants^54,61^. Further, TOR hyperactivation is associated with the downregulation of ABA signaling^22^. Therefore, we hypothesized that the FLZ8-mediated balancing of TOR-SnRK1 signaling would be linked with plants’ ability to respond to stress. To test this, we developed a dynamic network model of TOR-SnRK1 signaling in stress conditions (Fig. 7A). We simulated this network to understand how plants growing in favorable conditions respond to sudden exposure to stress with and without the FLZ8-mediated signaling network. The simulation predicted that FLZ8 helps in the rapid activation of SnRK1 and stress signaling leading to enhanced survival (Fig. 7B-D). To test these predictions, we first studied how SnRK1 signaling is regulated to sudden exposure to osmotic and salt stress. We estimated *DIN6* expression as it is widely used to monitor transient and physiologically relevant SnRK1 signaling^4,53–55^. Osmotic and salt stress treatments led to rapid induction of *DIN6* expression at 1h which was downregulated at 3h indicating a rapid activation of SnRK1 signaling under stress. This induction of *DIN6* expression at 1h was significantly attenuated in the *flz8* mutant (Fig. 7E and F). Thus, FLZ8 module which helps to maintain a basal level of SnRK1 activity under normal growth conditions helps in the rapid activation of SnRK1 signaling under stress.

**Fig. 7:**
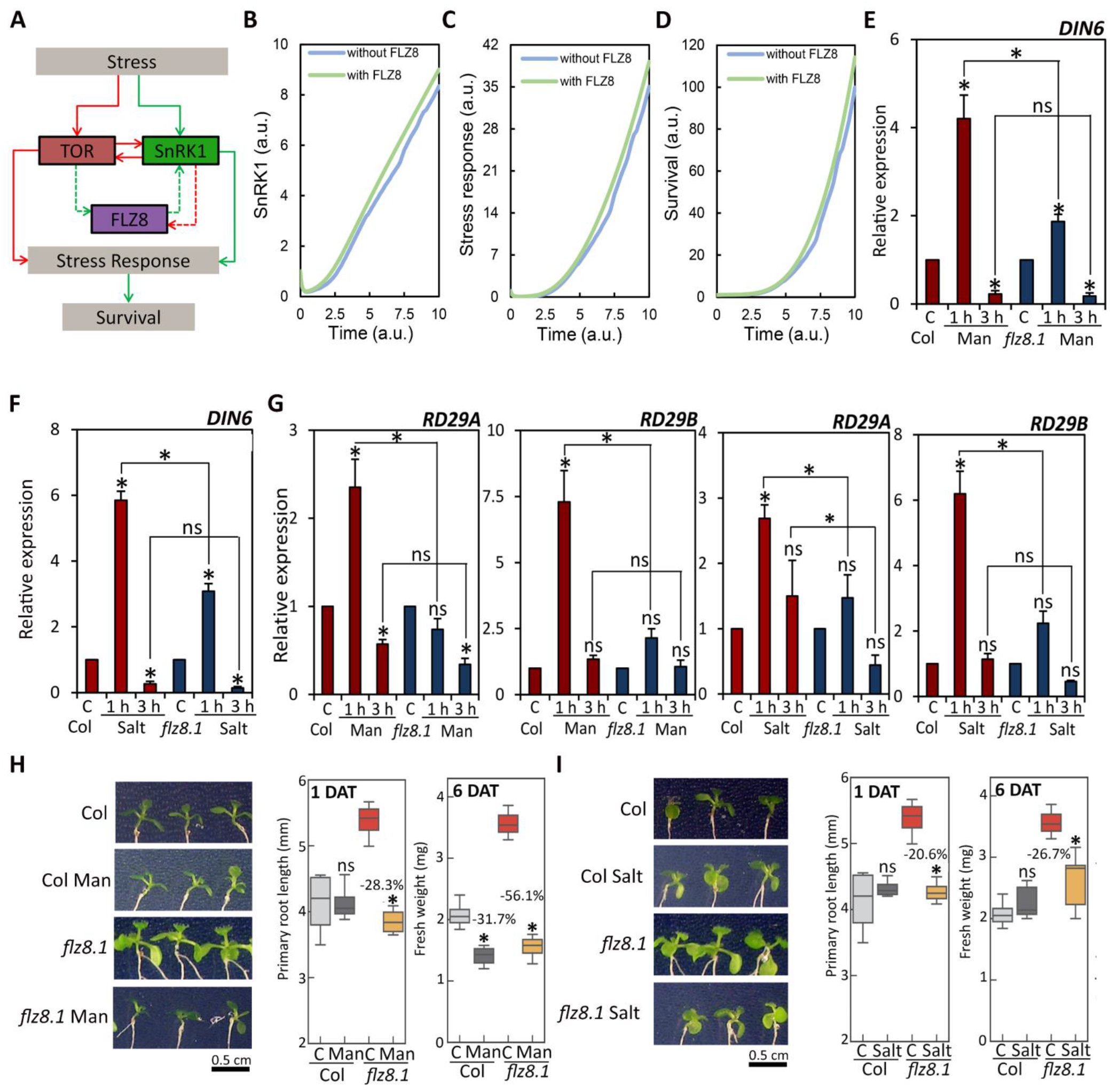
The FLZ8-mediated negative feedback promotes stress tolerance through the rapid activation of SnRK1 and stress signaling. (A) Network model developed for simulating the TOR-FLZ8-SnRK1 signaling controlling stress response. The green arrows indicate a positive influence and red arrows indicate a negative influence. The signaling networks which connect FLZ8 with other modules are represented by dotted arrows as these networks were absent while modeling the condition ‘without FLZ8’. (B)-(D) Simulation of SnRK1 activity, stress response activation and survival under sudden exposure to stress with and without FLZ8. (E) and (F) Induction of *DIN6* expression in response to short-term (1 and 3 h) osmotic (100 mM mannitol) and salt (50 mM NaCl) stresses in WT and *flz8.1* (two-way ANOVA, *p ≤ 0.05, Bonferroni post-hoc test). (G) Induction of *RD29A* and *RD29B* (abiotic stress marker genes) expression in response to short-term (1 and 3 h) osmotic (100 mM mannitol) and salt (50 mM NaCl) stress treatments in WT and *flz8.1* mutant (two-way ANOVA, *p ≤ 0.05, Bonferroni post-hoc test). (H) Effect of short-term (1 day) and long-term (6 days) osmotic stress (100 mM mannitol) treatments on root length and growth in WT and *flz8.1* (two-way ANOVA, *p ≤ 0.05, Bonferroni post-hoc test). (I) Effect of short-term (1 day) and long-term (6 days) salt stress (50 mM NaCl) treatments on root length and growth in WT and *flz8.1* (two-way ANOVA, *p ≤ 0.05, Bonferroni post-hoc test).

ABA and SnRK1 show interdependence in signaling activation and activate a large set of common genes^54,62,63^. Loss of *FLZ8* led to changes in TOR-SnRK1 signaling and ABA sensitivity indicating the perturbation in the ABA-mediated stress signaling. To test this, we estimated the induction of ABA-marker genes *Responsive to Desiccation 29A* (*RD29A*) and *Responsive to Desiccation 29B* (*RD29B*) in short-term osmotic and salt stress treatments in the *flz8* mutant. The stress treatments rapidly induced the expression of *RD29A* and *RD29B* at 1h which was found to be significantly attenuated in *flz8* mutant (Fig. 7G). Further, when exposed to osmotic and salt stress, the mutant showed more pronounced inhibition in root growth and fresh weight in comparison to the WT (Fig. 7H and I). Taken together, the expression and physiological analysis confirm the simulation results establishing the role of FLZ8 module for the timely activation of stress signaling machinery (Fig. S10).

## DISCUSSION

In this study, we identified a feedback regulatory mechanism involved in maintaining optimal level of TOR signaling in plants, which is critical for adjusting growth according to the environmental conditions. Once the TOR signaling is activated in response to sugar sufficiency^10,12^, how plants moderate TOR signaling to maintain signaling homeostasis is not understood. We report that FLZ8, a land plant-specific zinc finger protein^27,30^ is a critical regulator of TOR during sugar sufficiency operating in a negative feedback loop and this regulation was found to be crucial in moderating growth and rapid activation of stress signaling in plants (Fig. S10).

The reciprocal interaction of TOR and ABA signaling regulates growth and stress responses in plants^22^. However, this mutually antagonistic interaction operates either when plants are under nutrient sufficiency (i.e. high TOR activity leading to the suppression of ABA signaling) or stress (i.e. activation of ABA signaling leading to the suppression of TOR activity). In opisthokonts, TOR signaling is under tight negative feedback regulation in nutrient sufficiency^16,18^. As these regulatory proteins are absent in the plant lineage, we reasoned that plant might have evolved distinct regulatory mechanisms for adjusting their growth in the fluctuating environment on land. To test this, using the information on TOR-ABA signaling^22^, we modeled the potential role of negative feedback regulation of TOR and simulations identified its role in improving the fitness in a changing environment.

The high-throughput interaction and phosphoproteomics analyses identified the plant TOR signaling network in high resolution^28,64^. We integrated other experimental data with this to develop an updated network of TOR signaling in plants. Sugar level is one of the primary endogenous signals connected to TOR activity in plants^10,12^. Therefore, we screened the response of 265 TOR signaling network genes in Glc treatment to find potential negative feedback regulatory proteins. 7 genes were found to have strong correlation between transcriptional induction and Glc sufficiency. Among these genes, we focused our work on FLZ8 based on its interaction with both RAPTOR1B and SnRK1 subunits^27,28^. A paralog of FLZ8 named FLZ9/MARD1 also interacts with RAPTOR1B and SnRK1 subunits^28,30^. However, its expression was not induced by sugar and it is reported to be a primary ABA-inducible gene^31^.

Our detailed analysis identified the involvement of two distinct histone modifications in the induction of *FLZ8* in Glc sufficiency. The enrichment of H3KAc was found to be mediated through the TOR and HAC1 pathway and H3K4me3 enrichment was found to be independent of TOR signaling. Further, *FLZ8* expression was found to have a clear positive correlation with Suc and light availability. However, the expression of *FLZ8* didn’t show any positive correlation with the changes in the level of micronutrients which are also linked with TOR signaling in plants^45^. Thus, the sugar production through photosynthesis is a primary endogenous signal which enhances the *FLZ8* level through TOR-dependent and independent pathways. Further, SnRK1 activation under sugar starvation suppresses the level of *FLZ8*. Collectively, our results establish that the *FLZ8* level is highly sensitive to changes in the cellular sugar level mediated through TOR-SnRK1 signaling and other independent mechanisms.

Mutants of *FLZ8* showed TOR hyperactivity in favorable conditions which led to accelerated developmental responses such as the early opening of cotyledons, increased meristem size and activity. This led to enhanced growth and biomass in mutants. Strikingly, this phenotype was found to be strongly correlated with sugar status. The growth acceleration was more pronounced when shifted to high sugar (60 mM Suc) regime and the difference in growth was completely abolished when grown in sugar-limited (8mM Suc) regime. Taken together, these findings suggest that FLZ8 is a critical regulator of sugar-dependent growth through moderating signaling. The overlapping expression domain of *FLZ8* with TOR indicates that the TOR-FLZ8 module operates in the early seedling development to maintain the balance of TOR activity in plants.

We found that FLZ8 moderates TOR signaling through promoting SnRK1 signaling by bridging the RAPTOR1B-SnRK1α1 interaction and promoting SnRK1α1 stability. Studies in different eukaryotes identified that phosphorylation of RAPTOR by AMPK/SnRK1 is a conserved mechanism for the downregulation of TORC1 activity^5,65^. In plants, SnRK1α1 interacts and phosphorylates RAPTOR1B^66^. However, the molecular consequences of this phosphorylation in TOR signaling are yet to be identified. Nonetheless, phosphoproteomics analysis identified constitutive high phosphorylation of Ribosomal Protein S6, a major phosphorylation-target of TOR signaling in *snrk1α* mutant^66^. Thus, similar to other eukaryotes, the SnRK1α1 and RAPTOR1B interaction results in the phosphorylation of the latter and possibly leads to the downregulation of TOR activity. FLZ8 acts as a scaffold protein which promotes this association causing downregulation of TOR activity.

We found that FLZ8 interacts with the UBA domain of SnRK1α1. A recent report identified that UBA in SnRK1α1 promotes the phosphorylation of SnRK1 by activating kinases and helps in maintaining the kinase activity for longer duration^50^. In general, the UBA domain interacts with ubiquitin and prevents protein degradation^52^. In line with this, loss of *FLZ8* led to downregulation of the SnRK1α1 level. The reduction in the SnRK1α1 led to depleted level of SnRK1 active pool and reduced SnRK1 activity in *flz8* mutants. Thus, FLZ8 is involved in enhancing the stability of SnRK1α1 possibly by preventing ubiquitination leading to negative regulation of TOR.

The *FLZ* genes are part of the land-plants specific set of genes^27,30^. On land, plants are more exposed to adverse environmental conditions. The acquisition of novel genes and the evolution of new regulatory mechanisms was critical in the terrestrialization of plants^21^. Simulations identified that FLZ8 prevents hyperactivation of TOR and keeps a basal level of SnRK1 signaling during sugar sufficiency. Supporting this, loss of *FLZ8* led to pronounced growth in sugar-rich growth conditions. Further, we hypothesized that through this regulation, FLZ8 might be involved in the regulation of stress responses in plants. Simulations suggested that the FLZ8 module helps plants in the rapid activation of SnRK1 and stress signaling when exposed to stress. We tested this hypothesis using osmotic and salt stress conditions as SnRK1 and ABA signaling are involved in regulating stress responses under these conditions^4,22,67^. Analysis of the marker gene expression identified the role of FLZ8 in the rapid (1h) induction of SnRK1 and ABA signaling in stress. As a result, the mutant was hypersensitive to moderate osmotic and salt stress treatments. Thus, FLZ8 enables the cooption of TOR-SnRK1 signaling in plants to deal with the sudden changes in the environment (Fig. S9).

## METHODS

### Plant materials and growth conditions

The *Arabidopsis thaliana Columbia-0* (*Col-0*) ecotype was used for all experiments unless stated otherwise. The *SnRK1α1 OE2* line is in the Landsberg *erecta* (L*er*) background^4^. Therefore, L*er* was used as the wildtype control in the experiments with *SnRK1α1 OE2*. The *TOR OE1 (GK-166C06)*, *TOR OE2 (GK-548G07), tor-es1*, *tor 35-7, e2fa, hac1-2, hac1-3, hac1-6,* and *SnRK1α1 OE2* lines were described previously^4,10,34,35,68^. The T-DNA insertion lines of *FLZ8* (*flz8.1*: SALK_129608; *flz8.2*: SALK_137194C) obtained from ABRC were screened and homozygous lines were identified through PCR-based genotyping method. Insertion in *flz8.1* and *flz8.2* was found to be in the first exon after 675^th^ and 671^st^ nucleotide, respectively. The primers used are listed in Table S2.

The seeds were surface sterilized and stratified for 2 days in the dark at 4°C. The seeds were grown on 0.5X Murashige & Skoog medium (MS; pH: 5.7) with 30 mM Suc (Sigma-Aldrich) and 0.8% w/v agar (Himedia Laboratories, India) in square Petri plates. This composition was used for all experiments unless specified. The experiments on adult stages were performed in pots filled with agro peat and vermiculite (3:1). The plants were cultivated in a climate-controlled growth room under 16:8 h photoperiod at 22°C and a light intensity of 60 μmol m^−2^ sec^−1^.

### Data-driven screening

The genes which are involved in the TOR signaling network were retrieved from individual and high-throughput studies directly and through the protein-protein interaction repository BioGRID^69^. The data was manually curated for eliminating repeats and the refined network was visualized in Cytoscape (v3.7.2)^70^. The transcriptome data was retrieved from Genevestigator^71^ and heatmap visualization and hierarchical clustering (Pearson correlation algorithm with average linkage clustering) was performed in MultiExperiment Viewer (MeV, v4.8)^72^.

### Mathematical modeling

The regulatory networks are modeled using a set of ordinary differential equations (ODE’s). MATLAB’s ODE solver *ode45* was used to solve the Initial Value Problem (IVP) described in the equation. A detailed account of the regulatory network analysis, computational method and MATLAB codes are given as Supplementary methods.

### Phenotypic analyses

The *Col-0* and *flz8.1* and *flz8*.2 lines were grown on 0.5X MS medium with 30 mM Suc. The cotyledon angle was measured at 3 DAG. The seedlings were photographed with SteREO Discovery.V12 (Zeiss) stereo microscope and the angle was measured with ImageJ (NIH, USA). The meristem size and diameter were measured at 5 DAG. The seedlings were incubated for 2 min in clearing solution (chloral hydrate: water: glycerol in 8:3:1 ratio). The images were taken using the AxioImager M2 Imaging System (Zeiss) and meristem size and diameter were measured with ImageJ (NIH, USA). The fresh weight, primary root length and lateral root number were quantified at 10 DAG.

For the root growth kinetics assay, the seedlings were grown on 0.5X MS medium in standard growth conditions and root growth was marked every day from 3^rd^ day onwards for 10 days. The day-wise growth was measured with ImageJ (NIH, USA). For the phenotypic analysis at the adult stages, the *Col-0* and *flz8* mutant lines were directly grown on pots and photographs and biomass measurements were taken at 15, 20, 25 and 30 DAG.

### AZD8055 sensitivity assay

The 5 DAG *Col-0* and *flz8* seedlings were transferred to 0.5X MS medium with 0.25, 0.5, and 1 μM or without AZD8055. The seedlings were grown for another 5 days at the standard growth conditions and fresh weight was measured.

### ABA sensitivity assay

The 5 DAG *Col-0* and *flz8* seedlings were transferred to 0.5X MS medium with 0 and 5 μM ABA. The seedlings were grown for another 5 days at the standard growth conditions and root length was measured.

### Measurement of sugar-induced hypocotyl elongation in dark

The 5 DAG *Col-0* and *flz8* seedlings growing on 0.5X MS medium with 30 mM Suc were kept in darkness in standard growth conditions. The plates were photographed at 0, 1, 2, and 3 days of dark treatment and hypocotyl length was measured.

### Measurement of sugar-regulated biomass accumulation

The 5 DAG *Col-0*, *flz8*.1, and *tor 35-7* seedlings growing on 0.5X MS medium with 30 mM Suc were transferred to 0.5X MS medium with 60 mM Suc and grown at standard growth conditions for another 5 days. Simultaneously, a control experiment was performed in which seedlings grown on 0.5X MS medium with 30 mM Suc were transferred to 8 mM Suc.

### Abiotic stress treatments

The 5 DAG *Col-0* and *flz8* seedlings growing on 0.5X MS medium with 30 mM Suc were transferred to osmotic stress (0.5X MS, 100 mM mannitol, 30 mM Suc) and salt stress (0.5X MS, 50 mM NaCl, 30 mM Suc) medium. The root growth was measured using ImageJ (NIH, USA) and fresh weight was measured using a microbalance (Sartorius, Germany).

### Measurement of root meristem activity

Edu (5-ethynyl-2′-deoxyuridine) staining was performed using Click-iT® EdU AlexaFluor® 647 Imaging Kit as per the kit protocol (Invitrogen). The 5 DAG *Col-0* and *flz8* seedlings grown on 0.5X MS medium with 30 mM Suc were fixed in the fixation reagent (1X PBS with 3.7% formaldehyde and 0.5% Triton-X) for 30 min in a vacuum chamber. After fixing, the seedlings were washed three times in washing buffer (1X PBS with 3% BSA) and incubated in the reaction cocktail for 30 min in the dark. After the incubation, seedlings were washed with washing buffer followed by a wash in 1X PBS. The images were taken with AxioImager M2 Imaging System (Zeiss) (Excitation: 650 nm; Emission: 670 nm).

### GUS assay

The *pFLZ8::GUSA* line^32^ was used for the GUS assay. The GUS assay was performed as reported previously^73^ with 3 h incubation in GUS buffer. The photographs were taken with AxioImager M2 Imaging System (Zeiss) and SteREO Discovery.V12 (Zeiss) stereo microscopes.

### Chemical treatments and sample preparation for qRT-PCR

For Torin 1 treatment, the *Col-0* seedlings were grown on 0.5X MS solid medium with 30 mM Suc for 10 days. The 10 DAG seedlings were transferred to 0.5X MS and 30 mM Suc liquid medium with and without Torin 1 (10 μM) in a 6-well plate and incubated at 22°C with shaking (100 rpm) for 3 h and samples were frozen in liquid nitrogen and stored at −80°C. The *TOR OE* lines, *e2fa*, *flz8.1* and *flz8.2* were grown along with *Col-0* for 10 days on 0.5X MS solid medium and samples were harvested for qRT-PCR.

For the sugar starvation treatment, the L*er* and *SnRK1α1 OE2* line were grown for 5 days in 0.5X MS solid medium with 30 mM Suc. These seedlings were incubated for 24 h in dark in 0.5X MS liquid medium without Suc in a 6-well plate at 22°C with shaking (100 rpm). One batch of the sample was harvested after 24 h of starvation and used for qRT-PCR. The other batch was divided into two groups and in one group, the Suc-free medium was replaced by the medium with 30 mM Suc. Both Suc-free and Suc-replenished samples were harvested after 3 h of incubation at 22°C in dark with shaking at 100 rpm.

### RNA extraction and qRT-PCR analysis

The RNA isolation was performed by the TRIzol method using RNeasy Plant Mini Kit (Qiagen). The RNA quality and quantity were assessed by NanoDrop 2000 spectrophotometer (ThermoFisher Scientific). Equal quantity of RNA was used for cDNA preparation using the High-Capacity cDNA Reverse Transcription Kit (ThermoFisher Scientific). The qRT-PCR analysis was performed with 1:20 diluted cDNA samples using PowerUp SYBR Green Master Mix (ThermoFisher Scientific) on Step One Plus or ViiA 7 Real-Time PCR System (ThermoFisher Scientific). The primers used for qRT-PCR were designed with Primer Express v3.0 software (ThermoFisher Scientific). The primers used for qRT-PCR are listed in Table S2.

### Yeast two- and three-hybrid assays

The full-length and partial constructs were cloned in pGBKT7g (BD), pGADT7g (AD) and pYES-DEST52 vectors using Gateway cloning. The primers used for cloning are listed in Table S2. All experiments were performed in Y2HGold yeast strain as per the manufacturer’s protocol (Clontech). The plates and liquid cultures were grown in incubator at 30°C with agitation (200 rpm) for liquid cultures.

The transformation was performed using the EZ-Yeast transformation kit (MP Biomedicals). Before the experiments, the auto-activation and toxicity of the prepared BD constructs were tested by spotting an equal amount of yeast culture carrying different constructs on single dropout (SDO) plates supplemented with 5-Bromo-4-chloro-3-indoxyl-α-D-galactopyranoside (X-α-Gal; working concentration: 40 μg/ ml) (Gold Bio) alone or in combination with the yeast toxin Aureobasidin A (AbA; working concentration: 200 ng/ ml) (Clonetech). Along with AD and BD construct experiments, a negative control experiment with BD vector and AD construct was also performed. The positive colonies were selected on double drop out medium (DDO; -Trp, -Leu) and equal amount of cells were spotted on DDO (-Trp, -Leu) and Quadruple dropout (QDO; -Trp, -Leu, -His, -Ade) medium supplemented with AbA and X-α-Gal. The plates were incubated at 30°C for 3 days and photographed.

In Y3H experiments, the yeast cells containing AD and BD combinations were transformed with pYES-DEST52-FLZ8 or pYES-DEST52 and positive colonies were selected on DDO (-Leu, -Ura) and SDO (-Trp). The positive colonies were selected and grown in DDO (-Leu, -Ura) medium with Glc or Gal as the carbon source. The interaction was tested by spotting equal amounts of cells on DDO (-Leu, -Ura), QDO (-Trp, -Leu, -His, -Ade) and QDO medium with AbA and X-α-Gal (QDO/X/A). The plates were incubated at 30°C for 3 days and photographed.

### Protein extraction and western blotting

10 DAG seedlings were used for western blotting to detect the endogenous level of the phosphorylated (Phospho-p70 S6 kinase pThr389 antibody; dilution: 1:1000; catalog no.: MA5-15117, Invitrogen) and total (Ribosomal S6 kinase 1/2 antibody; dilution: 1:2000; catalog no.: AS12 1855, Agrisera) level of S6Ks, phosphorylated (Phospho-AMPKα (Thr172) Antibody; dilution: 1:1000; catalog no.: 2531, Cell Signaling Technology) and total (SNF1-related protein kinase catalytic subunit alpha KIN10 antibody; dilution: 1:2000; catalog no.: AS10 919, Agrisera) level of SnRK1α1, and ATG8 (Autophagy-related protein ATG8 antibody; dilution: 1:2000; catalog no.: AS14 2811, Agrisera). HSP90-2 antibody (Heat shock protein 90-2 antibody; dilution: 1:5000; catalog no.: AS11 1629, Agrisera) was used as the loading control. Total protein was isolated in ice-cold extraction buffer (137 mM NaCl, 2.7 mM KCl, 4.3 mM Na_2_HPO_4_, 1.47 mM KH_2_PO_4_, 10% v/v glycerol, 0.05% v/v Triton X-100) with 1/500 (v/v) plant-specific protease inhibitor cocktail and phosphatase inhibitor cocktail 3 (Sigma-Aldrich). The protein concentration was quantified by the Bradford method using Protein Assay Dye Reagent (Bio-Rad) in a POLARstar Omega plate reader (BMG Labtech). An equal amount of protein was boiled in Laemmli buffer for 10 min at 95°C and separated on 12.5% acrylamide gel and blotted on a nitrocellulose membrane (GE healthcare). Blocking was done with 0.5% w/v of BSA in Tris-buffered saline with 0.1% v/v Tween 20 (TBST). The blots were developed using Clarity Western ECL Substrate (Bio-Rad) and visualized by ChemiDoc XRS+ imaging system (Bio-Rad). After the first detection, the membranes were stripped using glycine buffer (200 mM glycine, 3.5 mM SDS, 0.1% v/v Tween 20, pH: 2.2). After washing three times with TBST, the stripped membranes were blocked and used for immunodetection with other antibodies as per the protocol detailed above. The blots were quantified by ImageJ (NIH, USA).

### SnRK1 activity assay

The total protein was isolated from equal amount of (≈ 1 g) 10 DAG seedlings of WT and *flz8* lines using ice-cold extraction buffer (50 mM Tris-HCl, pH 7.5, 150 mM NaCl, 1 mM EDTA, 0.05% Triton X-100) supplemented with 1/500 (v/v) plant-specific protease inhibitor cocktail and phosphatase inhibitor cocktail 3 (Sigma-Aldrich). 20 μL (bed volume) Pierce Protein A/G Agarose beads (ThermoFisher Scientific) were gently washed three times for 5 min each in phosphate buffered saline (PBS) and incubated with 1.5 μg anti-SnRK1α1 antibody (SNF1-related protein kinase catalytic subunit alpha KIN10 antibody; catalog no.: AS10 919, Agrisera) in 600 μL PBS on a rotator shaker for 1 h at room temperature with gentle shaking. The bead-antibody complex was gently washed three times in protein extraction buffer and incubated in 1 mg total protein for 3 h at 4°C on a rotator with gentle shaking. A negative control experiment with protein extraction buffer incubated with antibody-bead complex was also performed along with the test samples. After incubation, beads were gently washed three times using protein extraction buffer and reconstituted to 20 μL using the same buffer. From this, 15 μL beads were used for SnRK1 activity assay.

The SnRK1 activity assay was performed employing AMARA peptide as the substrate using the Universal Kinase Activity kit which works on the basis of the activity of a coupling phosphatase which removes the inorganic phosphate from the ADP produced due to kinase activity (R&D Systems). The experiment was conducted as per the manufacturer’s protocol. The OD was determined in a POLARstar Omega plate reader (BMG Labtech) and the relative SnRK1 activity in the mutants was determined with reference to WT. A positive control experiment in which ATP was replaced by ADP (1 mM) was also conducted.

### Chromatin immunoprecipitation assay

The ChIP assays were performed according to the previous protocol with minor modifications ^74^. The *Col-0* and *e2fa* mutant lines were grown for 7 days on 0.5X MS medium in standard growth conditions. 1 g tissue of each genotype was crosslinked through vacuum infiltration for 10 min using the 1% v/v formaldehyde solution supplemented with 0.4 M Suc, 10 mM Tris–HCl pH 8, 1 mM PMSF and 1 mM EDTA. The cross-linked samples were ground in liquid nitrogen and nuclei were isolated. The sonication was performed in ice cold water using Bioruptor Plus sonication device (40 sec on, 20 sec off for 20 cycles) (Diagenode). The DNA-protein complexes were immunoprecipitated using a serum containing anti-E2Fa antibodies and protein digestion and DNA precipitation were performed as per the previous protocol ^74^. The enrichment of target promoter regions was analyzed by qRT-PCR using the primers listed in Table S2.

To analyze sugar and TOR-dependent acetylation and methylation, the *tor-es1*^35^ line was grown for 7 days in 0.5X MS medium. On the 7^th^ day, seedlings were transferred to 0.5X MS medium supplemented with 10 μM estradiol or 10 μM DMSO. After 4 days, seedlings were used for sugar starvation for 24 h followed by Glc treatment (0 and 30 mM) for 3 h as described previously^75^. A small portion of the sample tissue was used for qRT-PCR analysis depicted in Fig. 1C. 1 g seedlings of each treatment were used for ChIP assay using the protocol described in the above paragraph. To analyze sugar and TOR-dependent acetylation and methylation, the *hac1* lines were grown for 7 days in 0.5X MS medium and sugar starvation and Glc treatment were performed as described previously^75^. The anti-H3K (9+14+18+23+27) acetylation (Abcam) and anti-H3K4me3 (Millipore) antibodies were used for immunoprecipitation to study acetylation and methylation, respectively. The enrichment of target promoter regions of *FLZ8* was analyzed by qRT-PCR using the primers listed in Table S2.

### Statistical analyses

Instant Clue^76^, SigmaPlot v13 Microsoft Excel 2010 were used for graph preparation and statistical analyses. All experiments were performed at least three times unless specified.

## DATA AVAILABILITY

The mutant/transgenic lines can be requested from the corresponding authors. The authors confirm that any other data related to the findings of this manuscript are available from corresponding authors upon request.

## CODE AVAILABILITY

The MATLAB code used for modeling is given in the ‘Supplementary methods’ section.

## ACKNOWLEDGMENTS

This work was supported by the Department of Biotechnology (Project Grant BT/PR8001/BRB/10/1211/2013 and NIPGR Core Grant to AL; DBT-Senior Research Fellowship to MS, NIPGR-Junior Research Fellowship to CTM), Department of Science and Technology (INSPIRE Faculty Programme Grant IFA18-LSPA110 to MJK) and University Grant Commission (UGC-Senior Research Fellowship to MS), Government of India. The authors acknowledge NIPGR Confocal Facility for their assistance and DBT-eLibrary Consortium (DeLCON) for providing access to e-resources. We thank Filip Rolland for providing *SnRK1α1 OE2* seeds and Dr. Christian Meyer for *tor 35-7* seeds. We thank Dr. Manoj Kumar and Dr. Pallavi Agarwal for insightful discussion and constructive criticism of the manuscript.

## AUTHOR CONTRIBUTIONS

MJK and AL conceived the study. MJK, SJ, and AL designed the experiments. MS performed ChIP-qPCR experiments. MS, MJK, and SJ performed physiological experiments. MJK and CTM performed the mutant screening. SS and MJK performed the computational analysis. MJK and SJ performed all other experiments. MJK wrote the paper. SJ, and AL reviewed the paper.

## COMPETING INTERESTS

The authors declare no competing interests

## MATERIALS & CORRESPONDENCE

The request for materials and any other correspondence should be addressed to Muhammed Jamsheer K or Ashverya Laxmi

## Notes

### Competing Interest Statement

The authors have declared no competing interest.

## REFERENCES

1. Jamsheer K, M., Jindal, S. & Laxmi, A. Evolution of TOR–SnRK dynamics in green plants and its integration with phytohormone signaling networks. J. Exp. Bot. 70, 2239–2259 (2019).

2. González, A. & Hall, M. N. Nutrient sensing and TOR signaling in yeast and mammals. EMBO J. 36, 397–408 (2017).

3. Lin, S.-C. & Hardie, D. G. AMPK: Sensing Glucose as well as Cellular Energy Status. Cell Metab. 27, 299–313 (2018).

4. Baena-González, E., Rolland, F., Thevelein, J. M. & Sheen, J. A central integrator of transcription networks in plant stress and energy signalling. Nature 448, 938–942 (2007).

5. Gwinn, D. M. et al. AMPK Phosphorylation of Raptor Mediates a Metabolic Checkpoint. Mol. Cell 30, 214–226 (2008).

6. Nukarinen, E. et al. Quantitative phosphoproteomics reveals the role of the AMPK plant ortholog SnRK1 as a metabolic master regulator under energy deprivation. Sci. Rep. 6, 31697 (2016).

7. Ling, N. X. Y. et al. mTORC1 directly inhibits AMPK to promote cell proliferation under nutrient stress. Nat. Metab. (2020). doi:10.1038/s42255-019-0157-1

8. Menand, B. et al. Expression and disruption of the Arabidopsis TOR (target of rapamycin) gene. Proc. Natl. Acad. Sci. U. S. A. 99, 6422–7 (2002).

9. Ren, M. et al. Target of Rapamycin Regulates Development and Ribosomal RNA Expression through Kinase Domain in Arabidopsis. PLANT Physiol. 155, 1367–1382 (2011).

10. Xiong, Y. et al. Glucose–TOR signalling reprograms the transcriptome and activates meristems. Nature 496, 181–186 (2013).

11. Pfeiffer, A. et al. Integration of light and metabolic signals for stem cell activation at the shoot apical meristem. Elife 5, (2016).

12. Li, X. et al. Differential TOR activation and cell proliferation in Arabidopsis root and shoot apexes. Proc. Natl. Acad. Sci. U. S. A. 114, 2765–2770 (2017).

13. Chen, G.-H., Liu, M.-J., Xiong, Y., Sheen, J. & Wu, S.-H. TOR and RPS6 transmit light signals to enhance protein translation in deetiolating Arabidopsis seedlings. Proc. Natl. Acad. Sci. U. S. A. 115, 12823–12828 (2018).

14. Zhang, Z. et al. TOR Signaling Promotes Accumulation of BZR1 to Balance Growth with Carbon Availability in Arabidopsis. Curr. Biol. 26, 1854–1860 (2016).

15. Beauchamp, E. M. & Platanias, L. C. The evolution of the TOR pathway and its role in cancer. Oncogene 32, 3923–3932 (2013).

16. Lee, J. H. et al. Sestrin as a feedback inhibitor of TOR that prevents age-related pathologies. Science 327, 1223–8 (2010).

17. Budanov, A. V. & Karin, M. p53 Target Genes Sestrin1 and Sestrin2 Connect Genotoxic Stress and mTOR Signaling. Cell 134, 451–460 (2008).

18. Jin, G. et al. Skp2-Mediated RagA Ubiquitination Elicits a Negative Feedback to Prevent Amino-Acid-Dependent mTORC1 Hyperactivation by Recruiting GATOR1. Mol. Cell 58, 989–1000 (2015).

19. Nedelcu, A. M. Comparative Genomics of Phylogenetically Diverse Unicellular Eukaryotes Provide New Insights into the Genetic Basis for the Evolution of the Programmed Cell Death Machinery. J. Mol. Evol. 68, 256–268 (2009).

20. Panchaud, N., Péli-Gulli, M.-P. & De Virgilio, C. SEACing the GAP that nEGOCiates TORC1 activation. Cell Cycle 12, 2948–2952 (2013).

21. de Vries, J. & Archibald, J. M. Plant evolution: landmarks on the path to terrestrial life. New Phytol. 217, 1428–1434 (2018).

22. Wang, P. et al. Reciprocal Regulation of the TOR Kinase and ABA Receptor Balances Plant Growth and Stress Response. Mol. Cell 69, 100–112.e6 (2018).

23. Nguyen, L. K. & Kholodenko, B. N. Feedback regulation in cell signalling: Lessons for cancer therapeutics. Semin. Cell Dev. Biol. 50, 85–94 (2016).

24. Brandman, O. & Meyer, T. Feedback loops shape cellular signals in space and time. Science 322, 390–395 (2008).

25. Muraro, D., Byrne, H. M., King, J. R. & Bennett, M. J. Mathematical Modelling Plant Signalling Networks. Math. Model. Nat. Phenom. 8, 5–24 (2013).

26. Jamsheer K, M. et al. FCS-like zinc finger 6 and 10 repress SnRK1 signalling in Arabidopsis. Plant J. 94, 232–245 (2018).

27. Jamsheer K, M. et al. The FCS-like zinc finger scaffold of the kinase SnRK1 is formed by the coordinated actions of the FLZ domain and intrinsically disordered regions. J. Biol. Chem. (2018). doi:10.1074/jbc.RA118.002073

28. Arabidopsis Interactome Mapping Consortium. Evidence for Network Evolution in an Arabidopsis Interactome Map. Science (80-.). 333, 601–607 (2011).

29. Foster, K. G. et al. Regulation of mTOR Complex 1 (mTORC1) by Raptor Ser ^863^ and Multisite Phosphorylation. J. Biol. Chem. 285, 80–94 (2010).

30. Jamsheer K, M., Mannully, C. T., Gopan, N. & Laxmi, A. Comprehensive Evolutionary and Expression Analysis of FCS-Like Zinc finger Gene Family Yields Insights into Their Origin, Expansion and Divergence. PLoS One 10, e0134328 (2015).

31. He, Y. & Gan, S. A novel zinc-finger protein with a proline-rich domain mediates ABA-regulated seed dormancy in Arabidopsis. Plant Mol. Biol. 54, 1–9 (2004).

32. Jamsheer K, M. & Laxmi, A. Expression of Arabidopsis FCS-Like Zinc finger genes is differentially regulated by sugars, cellular energy level, and abiotic stress. Front. Plant Sci. 6, 746 (2015).

33. Thoreen, C. C. et al. An ATP-competitive mammalian target of rapamycin inhibitor reveals rapamycin-resistant functions of mTORC1. J. Biol. Chem. 284, 8023–32 (2009).

34. Deprost, D. et al. The Arabidopsis TOR kinase links plant growth, yield, stress resistance and mRNA translation. EMBO Rep. 8, 864–870 (2007).

35. Xiong, Y. & Sheen, J. Rapamycin and glucose-target of rapamycin (TOR) protein signaling in plants. J. Biol. Chem. 287, 2836–42 (2012).

36. Laribee, R. N. Transcriptional and Epigenetic Regulation by the Mechanistic Target of Rapamycin Complex 1 Pathway. J. Mol. Biol. 430, 4874–4890 (2018).

37. Zhao, Y. & Garcia, B. A. Comprehensive Catalog of Currently Documented Histone Modifications. Cold Spring Harb. Perspect. Biol. 7, a025064 (2015).

38. Wan, W. et al. mTORC1 Phosphorylates Acetyltransferase p300 to Regulate Autophagy and Lipogenesis. Mol. Cell 68, 323–335.e6 (2017).

39. Chen, Q. et al. WRKY18 and WRKY53 coordinate with HISTONE ACETYLTRANSFERASE1 to regulate rapid responses to sugar. Plant Physiol. pp.00511.2019 (2019). doi:10.1104/pp.19.00511

40. Sharma, M., Banday, Z. Z., Shukla, B. N. & Laxmi, A. Glucose-Regulated HLP1 Acts as a Key Molecule in Governing Thermomemory. Plant Physiol. 180, 1081–1100 (2019).

41. Vandepoele, K. et al. Genome-wide identification of potential plant E2F target genes. Plant Physiology 139, 316–328 (2005).

42. Xiong, Y. et al. Glucose–TOR signalling reprograms the transcriptome and activates meristems. Nature 496, 181–186 (2013).

43. Dong, Y. et al. Sulfur availability regulates plant growth via glucose-TOR signaling. Nat. Commun. 8, 1174 (2017).

44. Couso, I. et al. Phosphorus Availability Regulates TORC1 Signaling via LST8 in Chlamydomonas. Plant Cell (2019). doi:10.1105/TPC.19.00179

45. Shi, L., Wu, Y. & Sheen, J. TOR signaling in plants: conservation and innovation. Development 145, dev160887 (2018).

46. Chan, T. F., Carvalho, J., Riles, L. & Zheng, X. F. A chemical genomics approach toward understanding the global functions of the target of rapamycin protein (TOR). Proc. Natl. Acad. Sci. U. S. A. 97, 13227–32 (2000).

47. Montané, M.-H. & Menand, B. TOR inhibitors: from mammalian outcomes to pharmacogenetics in plants and algae. J. Exp. Bot. 70, 2297–2312 (2019).

48. He, Y. et al. Networking Senescence-Regulating Pathways by Using Arabidopsis Enhancer Trap Lines. PLANT Physiol. 126, 707–716 (2001).

49. Nietzsche, M., Landgraf, R., Tohge, T. & B??rnke, F. A protein-protein interaction network linking the energy-sensor kinase SnRK1 to multiple signaling pathways in Arabidopsis thaliana. Curr. Plant Biol. 5, 36–44 (2016).

50. Emanuelle, S., Doblin, M. S., Gooley, P. R. & Gentry, M. S. The UBA domain of SnRK1 promotes activation and maintains catalytic activity. Biochem. Biophys. Res. Commun. 497, 127–132 (2018).

51. Kim, D.-H. et al. mTOR Interacts with Raptor to Form a Nutrient-Sensitive Complex that Signals to the Cell Growth Machinery. Cell 110, 163–175 (2002).

52. Heinen, C., Ács, K., Hoogstraten, D. & Dantuma, N. P. C-terminal UBA domains protect ubiquitin receptors by preventing initiation of protein degradation. Nat. Commun. 2, (2011).

53. Mair, A. et al. SnRK1-triggered switch of bZIP63 dimerization mediates the low-energy response in plants. Elife 4, (2015).

54. Rodrigues, A. et al. ABI1 and PP2CA phosphatases are negative regulators of Snf1-related protein kinase1 signaling in Arabidopsis. Plant Cell 25, 3871–84 (2013).

55. Ramon, M. et al. Default activation and nuclear translocation of the plant cellular energy sensor SnRK1 regulate metabolic stress responses and development. Plant Cell tpc.00500.2018 (2019). doi:10.1105/tpc.18.00500

56. Jossier, M. et al. SnRK1 (SNF1-related kinase 1) has a central role in sugar and ABA signalling in *Arabidopsis thaliana*. Plant J. 59, 316–328 (2009).

57. Liu, Y. & Bassham, D. C. TOR Is a Negative Regulator of Autophagy in Arabidopsis thaliana. PLoS One 5, e11883 (2010).

58. Soto-Burgos, J. & Bassham, D. C. SnRK1 activates autophagy via the TOR signaling pathway in Arabidopsis thaliana. PLoS One 12, e0182591 (2017).

59. Chen, L. et al. The AMP-Activated Protein Kinase KIN10 Is Involved in the Regulation of Autophagy inArabidopsis. Front. Plant Sci. 8, 1201 (2017).

60. Klionsky, D. J. et al. Guidelines for the use and interpretation of assays for monitoring autophagy (3rd edition). Autophagy 12, 1–222 (2016).

61. Margalha, L., Confraria, A. & Baena-González, E. SnRK1 and TOR: modulating growth-defense trade-offs in plant stress responses. J. Exp. Bot. 70, 2261–2274 (2019).

62. Gazzarrini, S., Tsuchiya, Y., Lumba, S., Okamoto, M. & McCourt, P. The Transcription Factor FUSCA3 Controls Developmental Timing in Arabidopsis through the Hormones Gibberellin and Abscisic Acid. Dev. Cell 7, 373–385 (2004).

63. Tsai, A. Y.-L. & Gazzarrini, S. AKIN10 and FUSCA3 interact to control lateral organ development and phase transitions in Arabidopsis. Plant J. 69, 809–821 (2012).

64. Van Leene, J. et al. Capturing the phosphorylation and protein interaction landscape of the plant TOR kinase. Nat. Plants 5, 316–327 (2019).

65. Hughes Hallett, J. E., Luo, X. & Capaldi, A. P. Snf1/AMPK promotes the formation of Kog1/raptor-bodies to increase the activation threshold of TORC1 in budding yeast. Elife 4, (2015).

66. Nukarinen, E. et al. Quantitative phosphoproteomics reveals the role of the AMPK plant ortholog SnRK1 as a metabolic master regulator under energy deprivation. Sci. Rep. 6, 31697 (2016).

67. Jamsheer K, M. et al. The FCS-LIKE ZINC FINGER 6 and 10 are involved in regulating osmotic stress responses in Arabidopsis. Plant Signal. Behav. 1–4 (2019). doi:10.1080/15592324.2019.1592535

68. Heisel, T. J., Li, C. Y., Grey, K. M. & Gibson, S. I. Mutations in HISTONE ACETYLTRANSFERASE1 affect sugar response and gene expression in Arabidopsis. Front. Plant Sci. 4, 245 (2013).

69. Oughtred, R. et al. The BioGRID interaction database: 2019 update. Nucleic Acids Res. 47, D529–D541 (2019).

70. Su, G., Morris, J. H., Demchak, B. & Bader, G. D. Biological Network Exploration with Cytoscape 3. Curr. Protoc. Bioinforma. 2014, 8.13.1–8.13.24 (2014).

71. Hruz, T. et al. Genevestigator V3: A Reference Expression Database for the Meta-Analysis of Transcriptomes. Adv. Bioinformatics 2008, 1–5 (2008).

72. Saeed, A. I. et al. [9] TM4 Microarray Software Suite. in Methods in enzymology 411, 134–193 (2006).

73. Jefferson, R. A., Kavanagh, T. A. & Bevan, M. W. GUS fusions: beta-glucuronidase as a sensitive and versatile gene fusion marker in higher plants. EMBO J. 6, 3901–7 (1987).

74. Saleh, A., Alvarez-Venegas, R. & Avramova, Z. An efficient chromatin immunoprecipitation (ChIP) protocol for studying histone modifications in Arabidopsis plants. Nat. Protoc. 3, 1018–1025 (2008).

75. Mishra, B. S., Singh, M., Aggrawal, P. & Laxmi, A. Glucose and Auxin Signaling Interaction in Controlling Arabidopsis thaliana Seedlings Root Growth and Development. PLoS One 4, e4502 (2009).

76. Nolte, H., MacVicar, T. D., Tellkamp, F. & Krüger, M. Instant Clue: A Software Suite for Interactive Data Visualization and Analysis. Sci. Rep. 8, 12648 (2018).

