## Supplementary Figures for "A negative feedback loop of the TOR signaling moderates growth and enables rapid sensing of stress signals in plants"

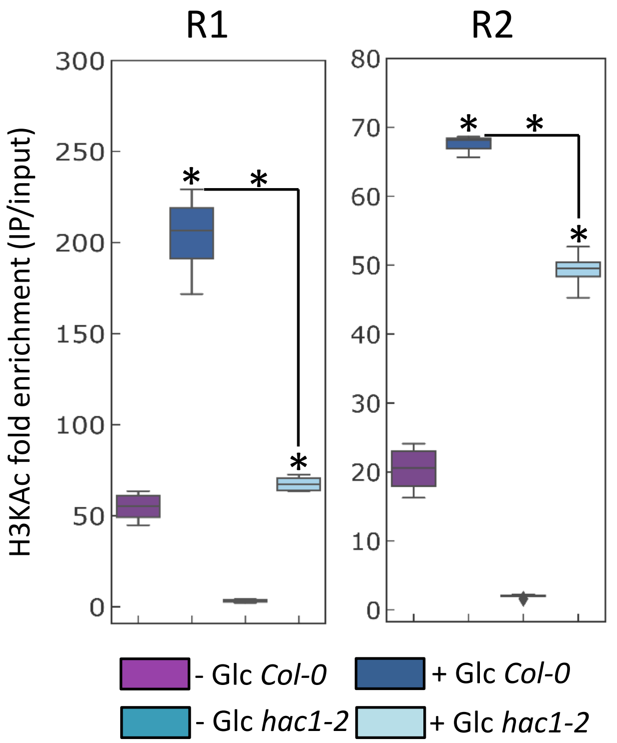


**Figure S1. HAC1 regulates the Glc-dependent H3KAc in the upstream regulatory regions of *FLZ8*.**
Histone acetylation (H3KAc) status in the upstream region of *FLZ8* promoter in *hac1-2* line treated with 0 or 170 mM Glc (two-way ANOVA, *p ≤ 0.05, Bonferroni post-hoc test).

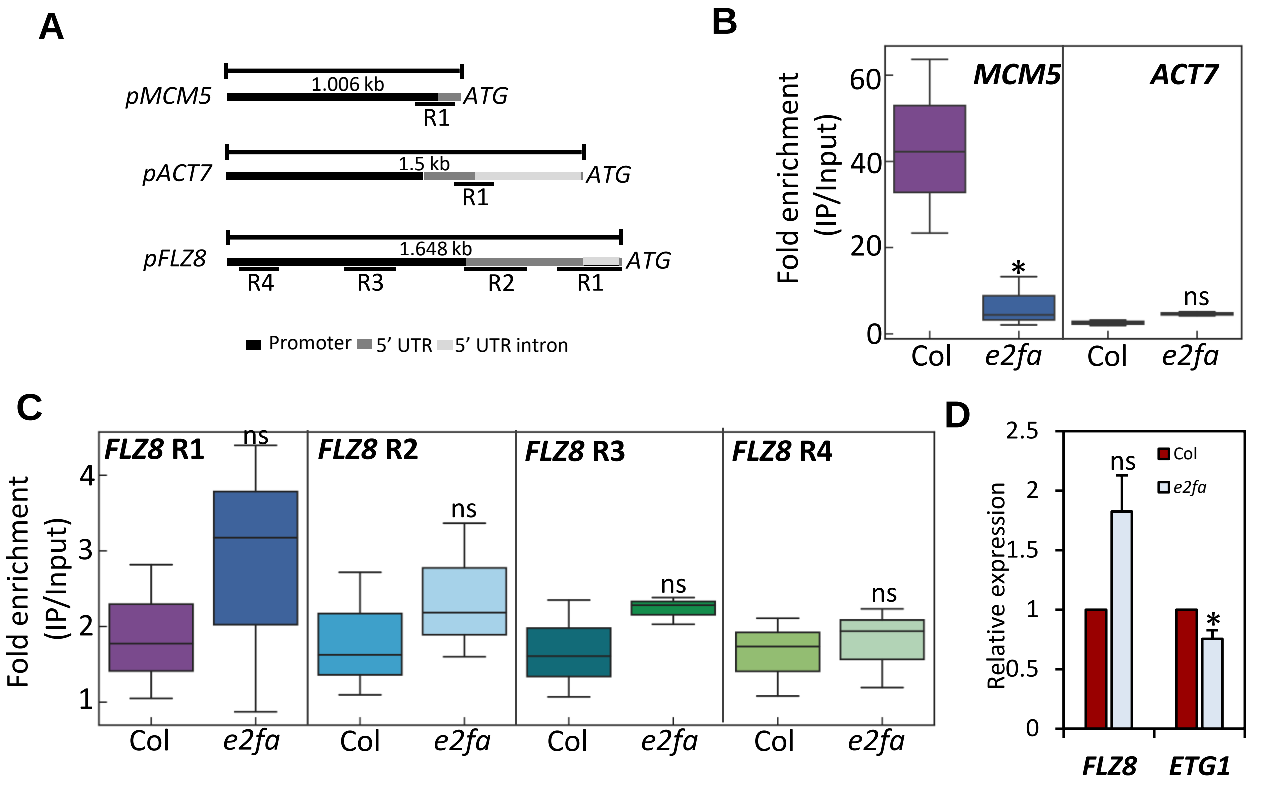


**Figure S2. Analysis of the role E2Fa in regulating the expression of *FLZ8*.**
(A) The promoter and other upstream regulatory regions of *MCM5*, *ATC7* and *FLZ8* indicating the regions tested for ChIP. (B) The binding of E2Fa to the upstream region of *MCM5* (positive control, a known target of E2Fa) and *ACT7* (negative control) in WT and *e2fa* mutant (one-way ANOVA, *p ≤ 0.05, Bonferroni post-hoc test). (C) The binding of E2Fa to the upstream regions of *FLZ8* in WT and *e2fa* mutant (one-way ANOVA, *p ≤ 0.05, Bonferroni post-hoc test). (D) Expression of *FLZ8* and *ETG1* (a known target of E2Fa) at 5DAG was quantified from *e2fa* in comparison to WT by qPCR analysis. The graph shown is the average of three biological replicates and error bar represents SE (one-way ANOVA, *p ≤ 0.05, Bonferroni post-hoc test).

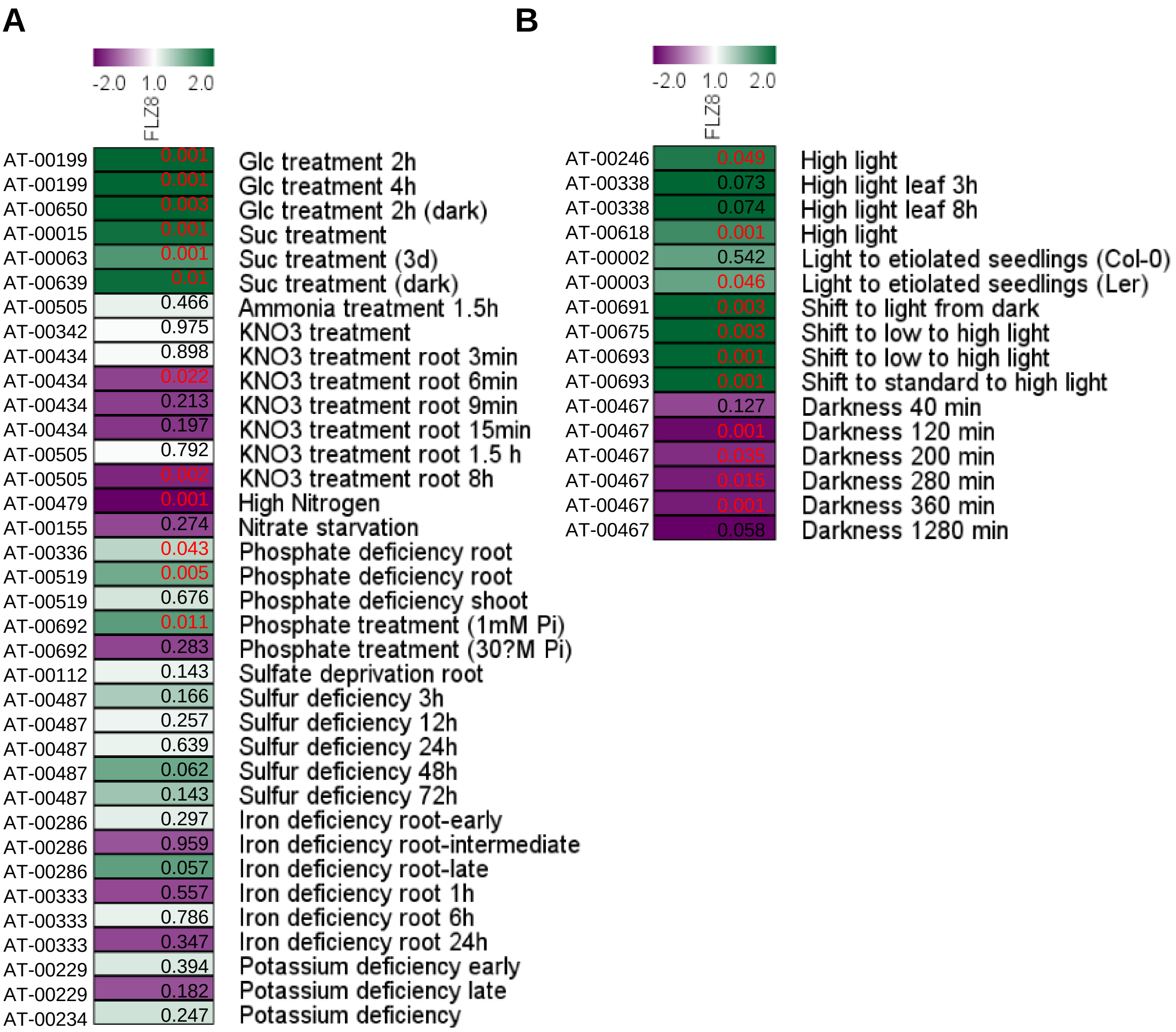


**Figure S3. Expression of *FLZ8* in different nutrient and light treatments.**

(A) Expression of *FLZ8* in different nutrient treatments. (B) Expression of *FLZ8* in different light conditions.

The expression of *FLZ8* in different nutrient and light treatments were analysed using Genevestigator. The value shown in the heat map is the p value showing statistical significance (red ≤ 0.05; black > 0.05).The code of transcriptome studies is given on the left side of the heatmap.

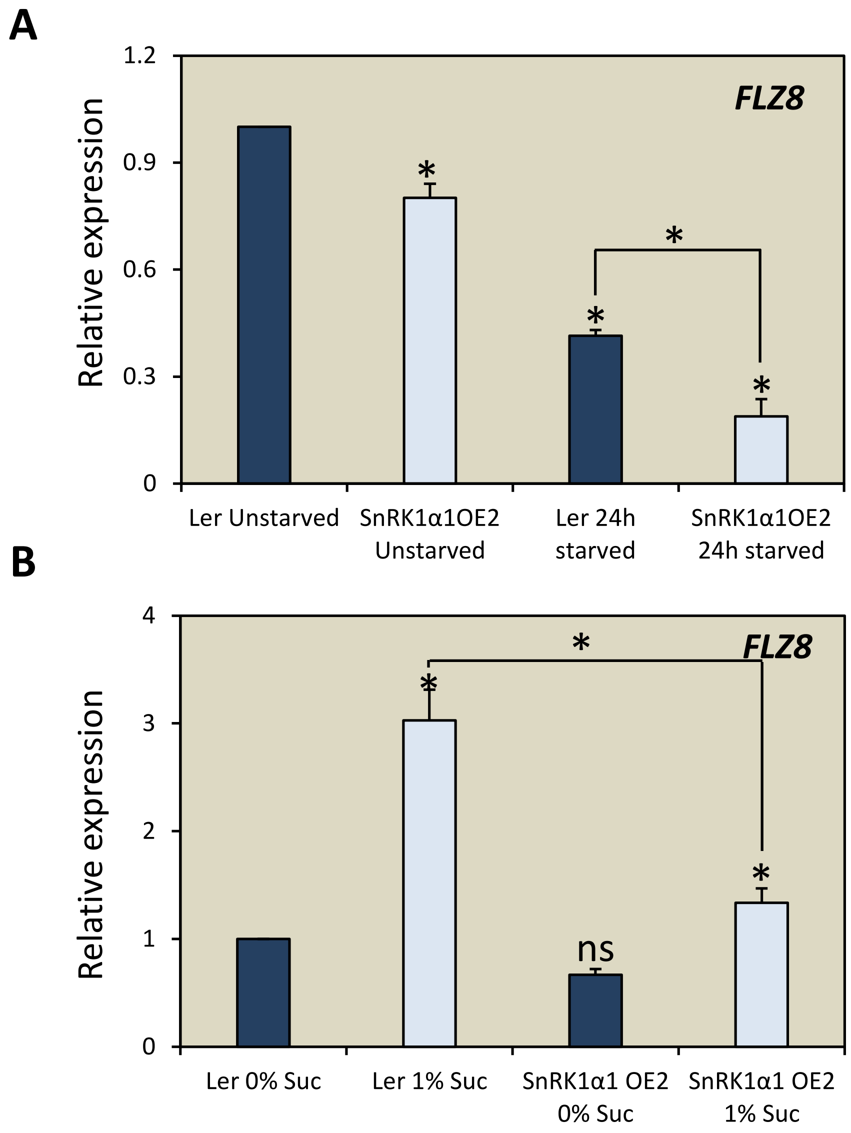


**Figure S4. SnRK1 signaling negatively regulates *FLZ8* transcript accumulation.**
(A) Transcript level of *FLZ8* during sucrose starvation in WT and *SnRK1α1 OE2* line in comparison to non-starved WT. (B) Transcript level of *FLZ8* in response to sucrose replenishment (30 mM in dark for 3 h) in WT and *SnRK1α1 OE2* line in comparison to WT grown in sucrose-starved condition (0 mM Suc & without light for 24 h). The graphs shown are the average of three biological replicates and error bar represents SE (two-way ANOVA, *$p \leq 0.05$, Bonferroni post-hoc test).

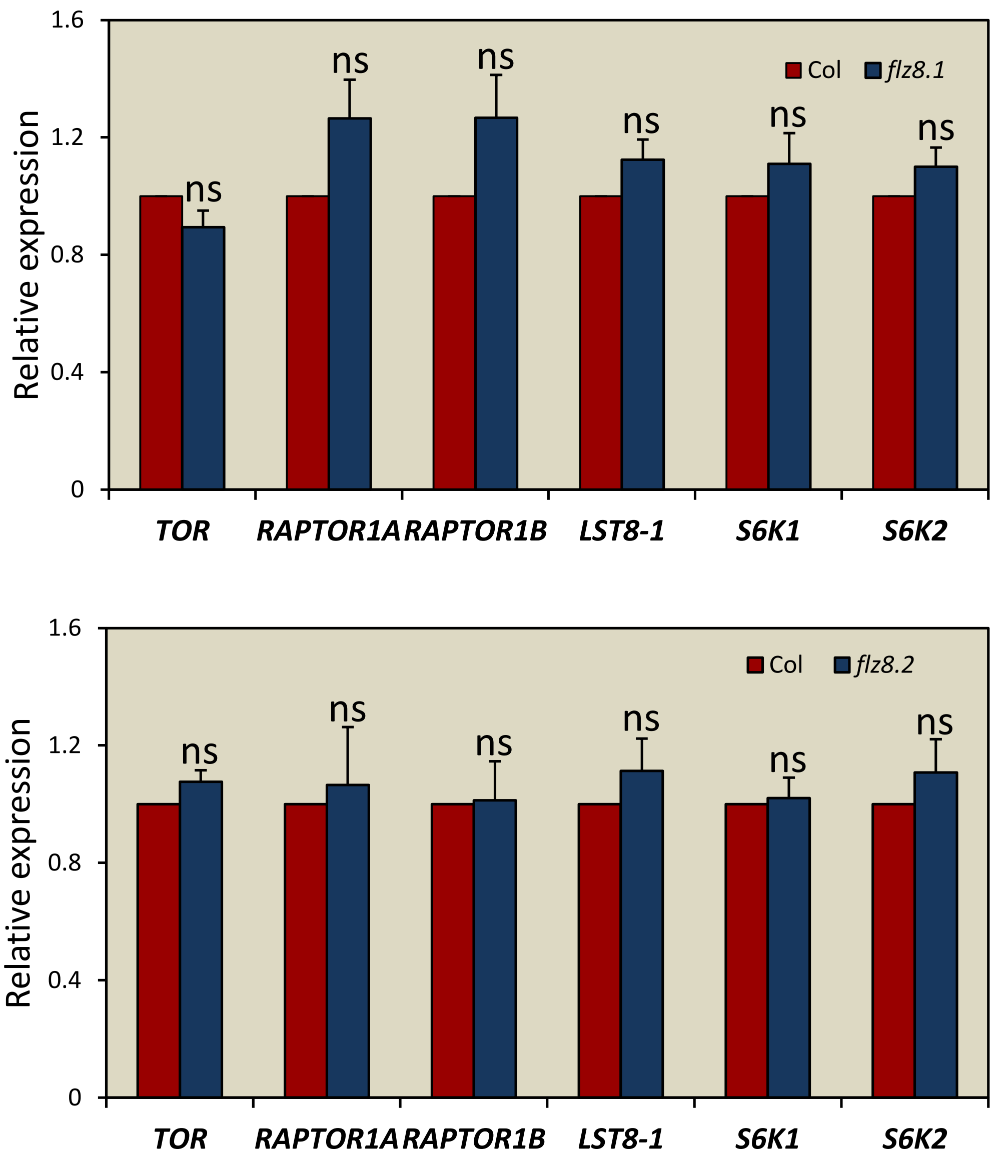


**Figure S5. Expression of TOR signaling related genes in *flz8* lines.**
Expression of TOR signaling related genes at 10 DAG was quantified from *flz8* mutant lines in comparison to WT by qPCR . The graphs shown are the average of three biological replicates and error bar represents SE (one-way ANOVA, *p ≤ 0.05, Bonferroni post-hoc test).

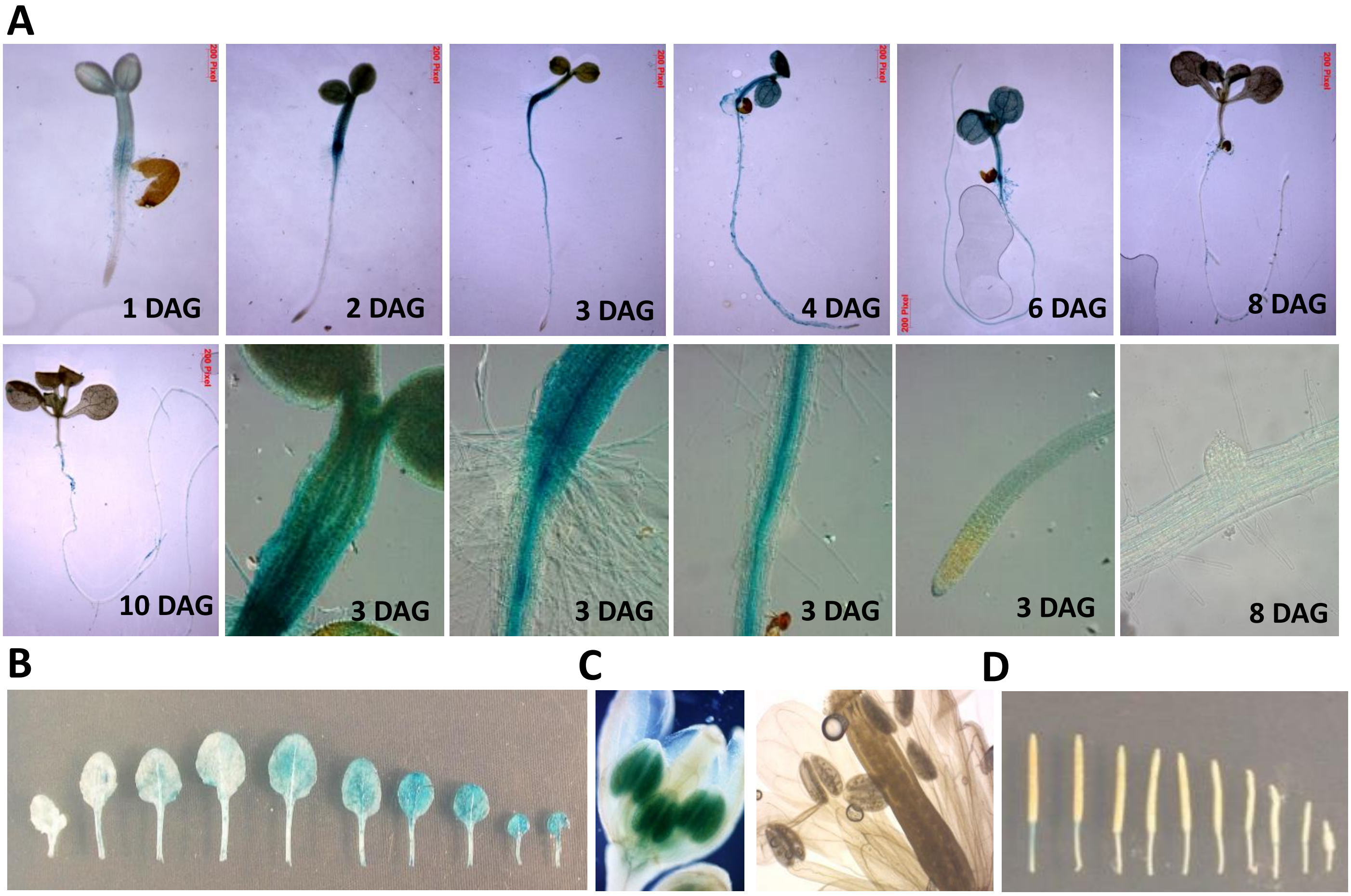


**Figure S6. Developmental stage and tissue-specific expression of *FLZ8*.**
(A) Activity of *pFLZ8::GUSA* in different stages of seedlings (DAG: Days After Germination). (B) Activity of *pFLZ8::GUSA* in leaves of 32 DAG rosette. The leaves were arranged according to increasing age from left to right. (C) Activity of *pFLZ8::GUSA* in flower bud and open flower. (D) Activity of *pFLZ8::GUSA* during silique development.

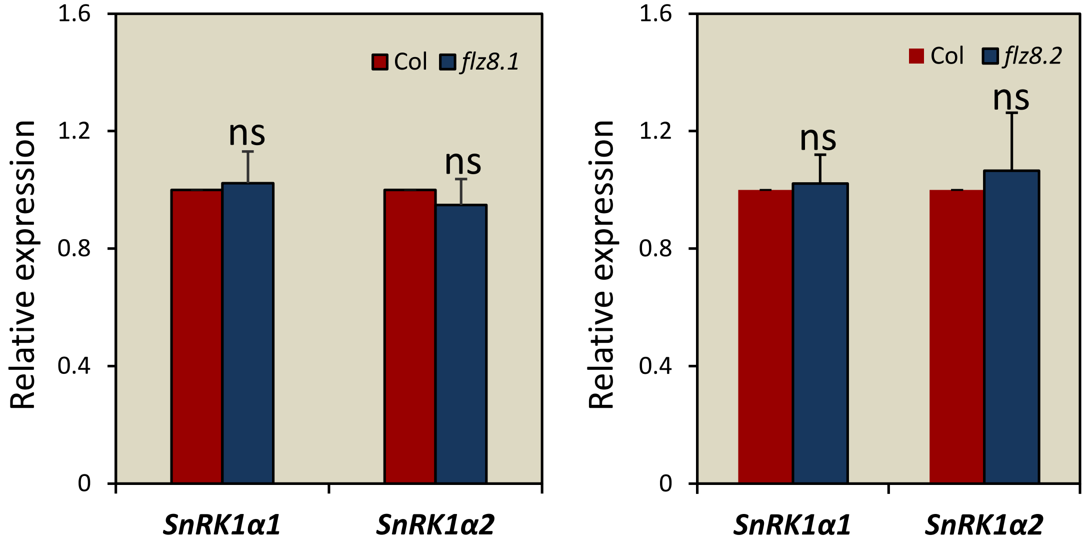


**Figure S7. Expression of *SnRK1α* genes in *flz8* lines.**
Expression of *SnRK1α* genes quantified from *flz8* mutant lines at 10 DAG in comparison to WT by qPCR analysis. The graph shown are the average of three biological replicates and error bar represents SE (one-way ANOVA, *$p \leq 0.05$, Bonferroni post-hoc test).

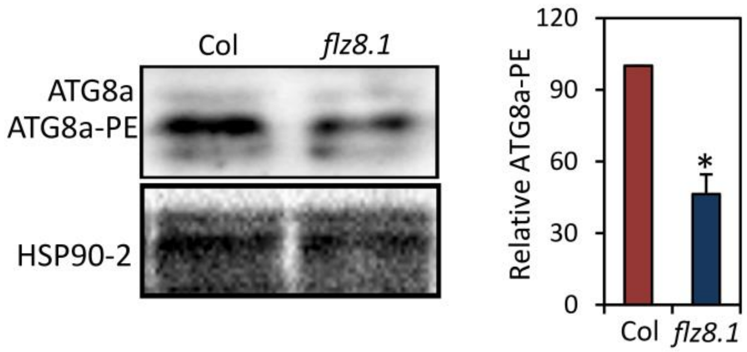


**Figure S8. Autophagy is negatively regulated in the *flz8* line.**
Level of ATG8a and its lipidated form ATG8a-PE in WT and *flz8.1*. The graphs represent the average of three biological replicates and error bars indicate SE (one-way ANOVA, *$p \leq 0.05$, Bonferroni post-hoc test).

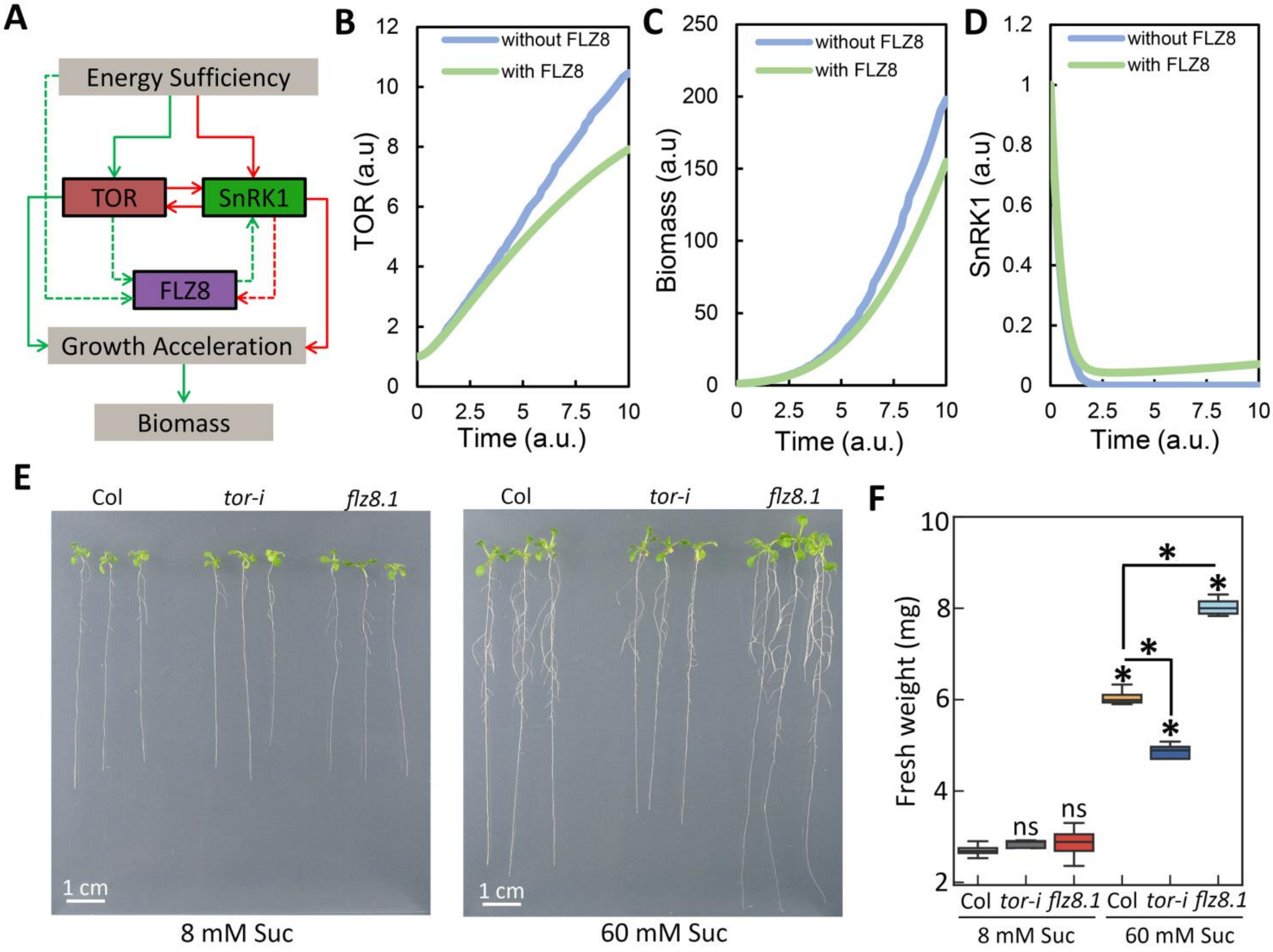


**Figure S9. The FLZ8 module negatively regulates TOR activity in the energy sufficiency to moderate growth.**

(A) Network model developed for simulating the TOR-FLZ8-SnRK1 signaling and growth in energy sufficiency. The green arrows indicate a positive influence and red arrows indicate a negative influence. The signaling networks which connect FLZ8 with other modules are represented by dotted arrows as these networks were absent while modeling the condition ‘without FLZ8’. (B)-(D) Simulation of TOR signaling, biomass accumulation and SnRK1 signaling under a favorable growth condition with and without FLZ8. (E) Phenotype of WT, *tor-i* (*tor 35-7*), and *flz8.1* in energy sufficiency (60 mM Suc) and deficiency (8 mM Suc). (F) Fresh weight in energy sufficiency (60 mM Suc) and deficiency (8 mM Suc) (two-way ANOVA, *p ≤ 0.05, Bonferroni post-hoc test).

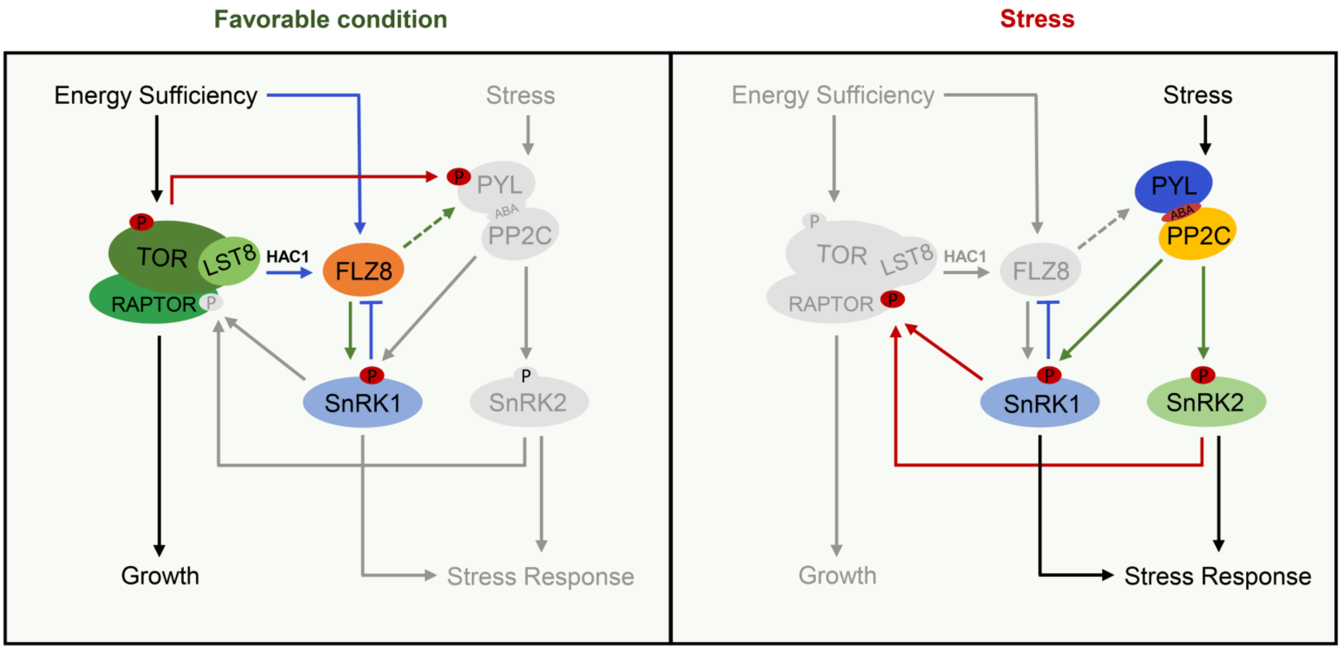


**Figure S10. Model illustrating the role of FLZ8-mediated negative feedback regulation of TOR signaling in controlling growth and stress response in plants.**

In favorable growth conditions, active TOR promotes growth and suppresses stress responses. At the same time, the level of *FLZ8* is upregulated by TOR and sugars through specific histone modifications. FLZ8 negatively regulates TOR signaling through enhancing the stability of SnRK1α1 and promoting the interaction of RAPTOR with SnRK1α1. Through this regulation, FLZ8 prevents TOR hyperactivation and maintains a basal level of SnRK1 and stress signaling which helps the plants to rapidly activate stress response when required. Key: Transcriptional regulations are depicted by blue arrows. The green arrows indicate positive regulation at the protein level (controlling protein stability and protein phosphorylation). Red arrows indicate protein phosphorylation events leading to a reduction in the activity of the substrate. The signaling events which are downregulated in each growth condition are shown in grey color.
