## Supplementary Methods for "A negative feedback loop of the TOR signaling moderates growth and enables rapid sensing of stress signals in plants"

### Mathematical modeling

To understand the significance of negative feedback regulation (denoted by X) in TOR signaling and growth and stress response, a regulatory network was modelled using a set of ordinary differential equations (ODE's). A regulatory network consists of interconnected nodes which represent the molecular activities.

$$\dot{y}_i = a_i + \sum_j p_{ji} y_j - k_i y_i - \sum_j d_{ji} y_j y_i \qquad (1)$$

In equation 1, $y_i$ denotes the activity of node i and $\dot{y}_i$ is the rate of activity, which is the time derivative. $a_i$ denotes the positive nodal activity at i, not explicitly related to the network under consideration (such as baseline protein production). $p_{ji}$ denotes the weight of the positive effect from node j to i, $d_{ji}$ denotes the weight of the negative effect from node j to i, both the weights being strictly positive. $k_i$ denotes the weight of negative effect not explicitly mentioned in the network under consideration (such as protein degradation).

The following list of symbols was used throughout the work. The $w_{ij}$ terms denote the weights in the network model.

The following list of symbols was used throughout the work. The $w_{ij}$ terms denote the weights in the network model.

$E \equiv Energy\ sufficiency$

$D \equiv Stress$

$T \equiv TOR\ signalling$

$X \equiv X\ factor\ signalling$

$A \equiv ABA\ signalling$

$S \equiv SnRK1\ signalling$

$F \equiv FLZ8\ signalling$

$G \equiv Growth\ acceleration$

$B \equiv Biomass$

$R \equiv Stress\ response$

$V \equiv Survival$

$w_{ET} \equiv Weight\ from\ node\ E\ to\ T$

$w_{AT} \equiv Weight\ from\ node\ A\ to\ T$

$w_{XT} \equiv Weight\ from\ node\ X\ to\ T$

$w_{TX} \equiv Weight\ from\ node\ T\ to\ X$

$w_{DA} \equiv Weight\ from\ node\ D\ to\ A$

$w_{TA} \equiv Weight\ from\ node\ T\ to\ A$

$w_{AV} \equiv Weight\ from\ node\ A\ to\ V$

$w_{ES} \equiv Weight\ from\ node\ E\ to\ S$

$w_{TS} \equiv Weight\ from\ node\ T\ to\ S$

$w_{ST} \equiv Weight\ from\ node\ S\ to\ T$

$w_{TF} \equiv Weight\ from\ node\ T\ to\ F$

$w_{SF} \equiv Weight\ from\ node\ S\ to\ F$

$w_{FS} \equiv Weight\ from\ node\ F\ to\ S$

$w_{EF} \equiv Weight\ from\ node\ E\ to\ F$

$w_{TG} \equiv Weight\ from\ node\ T\ to\ G$

$w_{SG} \equiv Weight\ from\ node\ S\ to\ G$

$w_{GB} \equiv Weight\ from\ node\ G\ to\ B$

$w_{DT} \equiv Weight\ from\ node\ D\ to\ T$

$w_{DS} \equiv Weight\ from\ node\ D\ to\ S$

$w_{TR} \equiv Weight\ from\ node\ T\ to\ R$

$w_{SR} \equiv Weight\ from\ node\ S\ to\ R$

$w_{RV} \equiv Weight\ from\ node\ R\ to\ V$

**Network model for TOR-X-ABA signaling**

In favourable growth conditions (unstressed), TOR phosphorylates PYL ABA receptors leading to suppression of ABA signaling. In stress conditions, ABA signaling activates SnRK2s which phosphorylate RAPTOR1B. This phosphorylation inhibits TOR-RAPTOR association and inhibition of TOR signaling[1]. Based on this antagonistic signaling interaction, a schematic model was developed depicting the interaction of these nodes with and without 'X' is given in Fig. 1a. The terms not mentioned explicitly in the schematic model (Fig. 1a) and presented in equation 1 were neglected. The set of ODE's formulated to represent the proposed regulatory network model are given in equation 2-8.

$$\dot{E} = 0 \quad (2)$$

$$\dot{T} = w_{ET}E - w_{AT}AT - w_{XT}XT \quad (3)$$

$$\dot{X} = w_{TX}T \quad (4)$$

$$\dot{G} = w_{TG}T \quad (5)$$

$$\dot{D} = 0 \quad (6)$$
$$\dot{A} = w_{DA}D - w_{TA}TA \quad (7)$$
$$\dot{V} = w_{AV}A \quad (8)$$

Equations 2-8 represent a system of coupled first order ODE's in seven variables. Normalized weights were used for quantifying the positive as well as negative effect between two nodes in the network path. The network model was also developed without 'X' by deleting the pathways connecting to 'X' (represented by dotted arrows in Fig. 1a) and thereby ignoring their weights.

We assumed normalized weights (0 to 1 in which 1 being the maximum influence) to all relations in the network. As 'X' is a negative feedback regulator, $\frac{1}{10}$ of the maximum weight was set as the maximum weight of all relations connected to 'X'.

**Network model for TOR-FLZ8-SnRK1 signaling in favourable conditions:** Our network consists of nodes which represent the effect of energy sufficiency condition on the activities of TOR, FLZ8 and SnRK1 to regulate growth acceleration and biomass. In eukaryotes including plants, TOR is activated and SnRK1 and its homologs (AMPK in mammals and SNF1 in yeast) are suppressed in response to energy sufficiency[2,3]. The positive correlation of TOR activity and negative correlation of SnRK1 activity with growth acceleration and biomass accumulation are also established[4–6]. SnRK1 through direct phosphorylation of TOR complex protein RAPTOR, and other specific mediator proteins (some of them are specific to certain eukaryotic lineages) suppress TOR function[2,7,8]. In yeast and mammals, TORC1 phosphorylates AMPK catalytic subunit leading to the suppression of AMPK activity[9]. Based on these previous studies and findings presented in this paper on the interaction of FLZ8 with TOR and SnRK1 signaling, a schematic model was developed depicting the interaction of these nodes which is given in Fig. S9a.

The system of equations was formulated using the similar approach used in the previous section. The set of ODE's formulated to represent the proposed regulatory network model are given in equations 9-14.

$$\dot{E} = 0 \quad (9)$$
$$\dot{T} = w_{ET}E - w_{ST}ST \quad (10)$$
$$\dot{S} = -w_{ES}ES - w_{TS}ST + w_{FS}F \quad (11)$$
$$\dot{F} = w_{TF}T + w_{EF}E - w_{SF}SF \quad (12)$$
$$\dot{G} = w_{TG}T - w_{SG}SG \quad (13)$$
$$\dot{B} = w_{GB}G \quad (14)$$

Equations 9-14 represent a system of coupled first order ODE's in six variables. Normalized weights were used for quantifying the positive as well as negative effect between two nodes in the network path. The network model was also developed without FLZ8 by deleting the pathways connecting to FLZ8 (represented by dotted arrows in Fig. S9a) and thereby ignoring their weights.

We assumed normalized weights (0 to 1 in which 1 being the maximum influence) to all relations in the network. In WT plants, sugar sufficiency linearly promotes growth and biomass accumulation in a TOR-dependent way indicating that although sugars and TOR promote FLZ8 in energy sufficiency condition, this pathway doesn't completely block sugar-dependent growth through SnRK1 activation. Thus, this pathway works as a fine-tuning mechanism to moderate growth in energy sufficiency. Therefore, compared to the normalized weights given to all other relations in the network, $\frac{1}{10}$ of the maximum weight was set as the maximum weight of all relations connected to FLZ8.

**Network model for TOR-FLZ8-SnRK1 signaling in unfavourable conditions:** In plants, TOR activity is reduced in response to several abiotic stress factors and TOR through direct phosphorylation of ABA receptors impinges ABA-mediated stress responses[1,10]. Further, TOR also suppresses Jasmonic Acid (JA) and (Salicylic Acid) (SA) pathways which make plants more susceptible to viral, bacterial and fungal attack[11]. ABA activates SnRK1 through the suppression of PP2C class of phosphatases which interact and dephosphorylate the Thr residue in the activation loop which is critical for their activity. SnRK1 and ABA signaling show a significant overlap in the transcriptional response[12]. SnRK1 also promotes JA and SA pathways to induce defence against viral, bacterial and fungal attack[13]. Thus, TOR and stress response machinery work antagonistically, while SnRK1 works largely synergistically with stress response machinery. Based on these previous studies and findings presented in this paper on the interaction of FLZ8 with TOR and SnRK1 signaling, a schematic model was developed depicting the interaction of these nodes which is given in Fig. 7a.

The system of equations was formulated using the similar approach used in the previous section. The set of ODE's formulated to represent the proposed regulatory network model are given in equations 15-20.

$$\dot{D} = 0 \tag{15}$$
$$\dot{T} = -w_{DT}DT - w_{ST}ST \tag{16}$$
$$\dot{S} = w_{DS}D - w_{TS}TS + w_{FS}F \tag{17}$$
$$\dot{F} = w_{TF}T - w_{SF}SF \tag{18}$$
$$\dot{R} = w_{SR}S - w_{TR}TR \tag{19}$$
$$\dot{V} = w_{RV}R \tag{20}$$

**Formulation of Initial value problem (IVP):**

The system of equations 2-8 is a system of seven first order equations. The seven dependant variables E, T, X, G, D, A and V are functions of the independent variable time (t), as the derivative involved is with respect to time. Integrating each equation leads to one arbitrary constant. For such system, infinite number of solutions exists. If the initial values of the variables are specified, then a unique solution can be obtained. So, the system requires seven initial conditions for a unique solution. An initial value problem (IVP) can be formulated using prescribed initial values of the variables. An IVP represents the dynamics of a system governed by a differential equation and depicts how the system evolves with time from a known initial state of the system. The equations 21-27 and equations 28-34 represent the IVP. The constants $E_0$, $T_0$, $X_0$, $G_0$, $D_0$, $A_0$ and $V_0$ are the prescribed initial values of the dependant variables in the proposed regulatory network.

To solve:

$$\dot{E} = 0 \tag{21}$$
$$\dot{T} = w_{ET}E - w_{AT}AT - w_{XT}XT \tag{22}$$
$$\dot{X} = w_{TX}T \tag{23}$$
$$\dot{G} = w_{TG}T \tag{24}$$

$$\dot{D} = 0 \quad (25)$$
$$\dot{A} = w_{DA}D - w_{TA}TA \quad (26)$$
$$\dot{V} = w_{AV}A \quad (27)$$

For the initial conditions:

$$E(at\ t = 0) = E_0 \quad (28)$$
$$T(at\ t = 0) = T_0 \quad (29)$$
$$X(at\ t = 0) = X_0 \quad (30)$$
$$G(at\ t = 0) = G_0 \quad (31)$$
$$D(at\ t = 0) = D_0 \quad (32)$$
$$A(at\ t = 0) = A_0 \quad (33)$$
$$V(at\ t = 0) = V_0 \quad (34)$$

In a similar fashion, the IVP for the TOR-FLZ8-SnRK1 signaling network for favourable conditions equations 9-14 can be formed as:

To solve:

$$\dot{E} = 0 \quad (35)$$
$$\dot{T} = w_{ET}E - w_{ST}ST \quad (36)$$
$$\dot{S} = -w_{ES}ES - w_{TS}ST + w_{FS}F \quad (37)$$
$$\dot{F} = w_{TF}T + w_{EF}E - w_{SF}SF \quad (38)$$
$$\dot{G} = w_{TG}T - w_{SG}SG \quad (39)$$
$$\dot{B} = w_{GB}G \quad (40)$$

For the initial conditions:

$$E(at\ t = 0) = E_0 \quad (41)$$
$$T(at\ t = 0) = T_0 \quad (42)$$
$$S(at\ t = 0) = S_0 \quad (43)$$
$$F(at\ t = 0) = F_0 \quad (44)$$
$$G(at\ t = 0) = G_0 \quad (45)$$
$$B(at\ t = 0) = B_0 \quad (46)$$

Similarly, the IVP for the TOR-FLZ8-SnRK1 signaling network for unfavourable conditions equations 15-20 can be formed as:

$$\dot{D} = 0 \quad (47)$$
$$\dot{T} = -w_{DT}DT - w_{ST}ST \quad (48)$$
$$\dot{S} = w_{DS}D - w_{TS}TS + w_{FS}F \quad (49)$$
$$\dot{F} = w_{TF}T - w_{SF}SF \quad (50)$$

$$\dot{R} = w_{SR}S - w_{TR}TR \quad (51)$$
$$\dot{V} = w_{RV}R \quad (52)$$

For the initial conditions:

$$D(at\ t = 0) = D_0 \quad (53)$$
$$T(at\ t = 0) = T_0 \quad (54)$$
$$S(at\ t = 0) = S_0 \quad (55)$$
$$F(at\ t = 0) = F_0 \quad (56)$$
$$R(at\ t = 0) = R_0 \quad (57)$$
$$V(at\ t = 0) = V_0 \quad (58)$$

Where, $D_0$, $T_0$, $S_0$, $F_0$, $R_0$, $V_0$ denote the prescribed initial values of the dependant variables in the proposed regulatory network.

**Computation:**

The IVP described by equations 21-27 and equations 28-34 was solved using MATLAB's ODE solver *ode45*. The solver uses numerical integration scheme known as Runge-Kutta (4-5) method, which is a combination of 4th and 5th order Runge-Kutta methods. The function *ode45* requires three basic inputs from the user. The solver integrates only the first order equations. In case of higher order differential equations, they must be converted into a system of first order equations. Consider an IVP as an illustrative example: $\dot{y} = f(y, t)$, with an initial value y(at t=$t_o$) =$y_0$, where $t_o$ is the initial time. The typical usage of *ode45* is illustrated below.

```
[t,y]=ode45(@DerivativeFunction,TimeSpan,InitialValue)
```

Note that "DerivativeFunction", "TimeSpan" and "InitialValue" are the identifiers used in the program; any other desired names which are permissible by MATLAB, can also be used by the reader.

Since the solver uses numerical integration for solving the IVP, it requires time-step. The solution will be computed for each time point based on the time-step. *ode45* uses a variable timestep for integration. The solver routines evaluate the timesteps based on the integration tolerances. In numerical integration, a final time should be prescribed to stop the integration. "TimeSpan" is an array consisting of the initial time ($t_o$) and the final time ($t_f$).

TimeSpan=[$t_o$ $t_f$];

"DerivativeFunction" is a user defined function which returns the derivatives from the differential equations, computed for each timestep, to be used by the solver. The function must be defined at the end of the program. The syntax for defining a user defined function in MATLAB is given below:

```
function dy=DerivativeFunction(t,y)

… The derivatives from the differential equations are defined here…

… dy is the array of derivatives…

end
```

The array “InitialValue” should contain the initial values of the dependant variables. For the IVP used in the illustration, the initial value of the dependant variable y is $y_0$. This condition is expressed in MATLAB as:

InitialValue= [$y_0$];

The MATLAB codes for the proposed regulatory networks are explained in the upcoming sections.

**TOR-X-ABA signaling in favourable and unfavourable conditions:**

To predict the role of ‘X’ module in growth in favourable and unfavourable conditions, we assumed an equal initial value (1 unit each) for TOR and ABA as base. Using the network model described above; we predicted the growth with and without ‘X’ under energy sufficiency (1 unit). The outcome was presented in Fig.1c and d.

**1. With X**

The equations 21-27 are in the first order form. The array of seven unknowns (E, T, X, G, D, A and V) were assumed as y=[y(1) y(2) y(3) y(4) y(5) y(6) y(7)]. Note that the order of the unknowns does not matter. An array of the seven derivatives was assumed accordingly, as dy. Consequently, the system of equations 21-27 were rewritten as:

$$dy(1) = 0 \tag{59}$$
$$dy(2) = w_{ET}y(1) - w_{AT}y(6)y(2) - w_{XT}y(3)y(2) \tag{60}$$
$$dy(3) = w_{TX}y(2) \tag{61}$$
$$dy(4) = w_{TG}y(2) \tag{62}$$
$$dy(5) = 0 \tag{63}$$
$$dy(6) = w_{DA}y(5) - w_{TA}y(2)y(6) \tag{64}$$
$$dy(7) = w_{AV}y(6) \tag{65}$$

A simple MATLAB code with all the descriptions is given below. Note that the symbol “%” should be used in the code for user comments.

```
%% MATLAB code for the solution for favourable and favourable condition with ‘X’
% The MATLAB solver function ode45 requires three basic inputs
% 1. The derivatives as per the system of equations (vector)
% 2. Initial values of the problem (vector)
% 3. The time span or domain of the problem (vector)
% MATLAB decides the time points from the initial and the final time
% using the timestep.
% Here, the variables were assumed in the order y=(E, T, X, G, D, A and V)
% “ode45” returns all the solutions at each time point.

TimeSpan=[0 10]; % Initial time is 0 and the final time is 10
InitialValue=[1 1 0 0 0 1 0]; % Prescribed initial values for E, T, X, G, D, A and V based on
%equations 59-65
%% Main function
% [y] in [t,y] holds all the solutions (E, T, X, G, D, A and V) at each time point.
% [y] is an array of 7 columns and as many as rows for the number of time steps
% [t] in [y,t] contains all the time points.
```

```
[t,y]=ode45(@ode_seven,TimeSpan,InitialValue); % Ode45 solver

%% Sample plot for the solution
% Plot for G (Growth), which is the 4th unknown
plot(t,y(:,4)); % Plotting the 4th column of the matrix [y] in Y-axis
% with time (t) on the X-axis.

%%%%%%%%%%%%%%%%%%%%%%%%%%%%%%%%%%%%%%%%%%%%%%%%%%%%%%%%%%%%%%%%%%%%
% Write here your code to plot any other solution


%%%%%%%%%%%%%%%%%%%%%%%%%%%%%%%%%%%%%%%%%%%%%%%%%%%%%%%%%%%%%%%%%%%%

function dy = ode_seven(t,y) % Function which returns the derivatives
% Assumed weights are the following
w_ET=1; w_AT=1;
w_TX=0.1; w_XT=0.1;
w_TG=1; w_DA=1; w_TA=1; w_AV=1;
dy=zeros(6,1); % Array initialization to all zeros
% Equations from 59-65
dy(1)=0;
dy(2)=w_ET*y(1)-w_AT*y(6)*y(2)-w_XT*y(3)*y(2);
dy(3)=w_TX*y(2);
dy(4)=w_TG*y(2);
dy(5)=0;
dy(6)=w_DA*y(5)-w_TA*y(2)*y(6);
dy(7)=w_AV*y(6);
end
```

**2. Without X**

In this case, the number of unknowns in the equations 21-27 was reduced to six as the variable X was not considered. The system consists of all the first order equations. The variables (E, T, G, D, A and V) were assumed as y=[y(1) y(2) y(3) y(4) y(5) y(6)]. The pathways consisting of X were deleted, which caused the corresponding weights to be set to zero. dy reduced to an array of six derivatives. Consequently, the system of equations became:

| | | |
|---|---|---|
| | $dy(1) = 0$ | (66) |
| | $dy(2) = w_{ET}y(1) - w_{AT}y(6)y(2)$ | (67) |
| | $dy(3) = w_{TG}y(2)$ | (68) |
| | $dy(4) = 0$ | (69) |
| | $dy(5) = w_{DA}y(4) - w_{TA}y(2)y(5)$ | (70) |
| | $dy(6) = w_{AV}y(5)$ | (71) |

The MATLAB code is given below. Some of the comments are omitted for compactness.

```
%% MATLAB code for the solution for favourable and favourable condition without 'X'
```

```
TimeSpan=[0 10]; % Initial time is 0 and the final time is 10
InitialValue=[1 1 0 0 1 0]; % Prescribed initial values for E, T, G, D, A and V based on
%equations 66-71
%% Main function
 [t,y]=ode45(@ode_six,TimeSpan,InitialValue); % Ode45 solver

%% Sample plot for the solution
% Plot for G (Growth), which is the 3rd unknown
plot(t,y(:,3));

%%%%%%%%%%%%%%%%%%%%%%%%%%%%%%%%%%%%%%%%%%%%%%%
%%%%%%%%%%%%%%%%%%%%%
% Write here your code to plot any other solution


%%%%%%%%%%%%%%%%%%%%%%%%%%%%%%%%%%%%%%%%%%%%%%%
%%%%%%%%%%%%%%%%%%%%%

function dy = ode_six(t,y) % Function which returns the derivatives
% Assumed weights are the following
w_ET=1; w_AT=1;
w_TG=1; w_DA=1; w_TA=1; w_AV=1;
dy=zeros(6,1); % Array initialization to all zeros
% Equations from 66-71
dy(1)=0;
dy(2)=w_ET*y(1)-w_AT*y(5)*y(2);
dy(3)=w_TG*y(2);
dy(4)=0;
dy(5)=w_DA*y(4)-w_TA*y(2)*y(5);
dy(6)=w_AV*y(5);
end
```

**TOR-X-ABA signaling in unfavourable condition**

**1. With X**

The equations 59-65 were used. The initial conditions were modified for the equations. The system assumed to have stress (1 unit) and the initial value of the TOR was assumed to be 10 units with baseline ABA signaling (1 unit). The MATLAB code is given below.

```
%% MATLAB code for the solution for favourable and unfavourable condition with 'X'
TimeSpan=[0 10]; % Initial time is 0 and the final time is 10
InitialValue=[0 10 0 0 1 1 0]; % Prescribed initial values for E, T, X, G, D, A and V based on
%equations 59-65
%% Main function
 [t,y]=ode45(@ode_seven,TimeSpan,InitialValue); % Ode45 solver

%% Sample plot for the solution
% Plot for G (Growth), which is the 4th unknown
plot(t,y(:,4));
```

```
%%%%%%%%%%%%%%%%%%%%%%%%%%%%%%%%%%%%%%%%%%%%%
%%%%%%%%%%%%%%%%%%%%
% Write here your code to plot any other solution


%%%%%%%%%%%%%%%%%%%%%%%%%%%%%%%%%%%%%%%%%%%%%
%%%%%%%%%%%%%%%%%%%%

function dy = ode_seven(t,y) % Function which returns the derivatives
% Assumed weights are the following
w_ET=1; w_AT=1;
w_TX=0.1; w_XT=0.1;
w_TG=1; w_DA=1; w_TA=1; w_AV=1;
dy=zeros(6,1); % Array initialization to all zeros
% Equations from 59-65
dy(1)=0;
dy(2)=w_ET*y(1)-w_AT*y(6)*y(2)-w_XT*y(3)*y(2);
dy(3)=w_TX*y(2);
dy(4)=w_TG*y(2);
dy(5)=0;
dy(6)=w_DA*y(5)-w_TA*y(2)*y(6);
dy(7)=w_AV*y(6);
end
```

**2. Without X**

The equations 66-71 were used. The same initial conditions as in the previous section were considered. The MATLAB code is given below.

```
%% MATLAB code for the solution for favourable and unfavourable condition with 'X'
TimeSpan=[0 10]; % Initial time is 0 and the final time is 10
InitialValue=[0 10 0 1 1 0]; % Prescribed initial values for E, T, G, D, A and V based on
%equations 66-71
%% Main function
 [t,y]=ode45(@ode_six,TimeSpan,InitialValue); % Ode45 solver

%% Sample plot for the solution
% Plot for G (Growth), which is the 3rd unknown
plot(t,y(:,3)); % Plotting the 3rd column of the matrix [y] in Y-axis
% with time (t) on the X-axis.

%%%%%%%%%%%%%%%%%%%%%%%%%%%%%%%%%%%%%%%%%%%%%
%%%%%%%%%%%%%%%%%%%%
% Write here your code to plot any other solution


%%%%%%%%%%%%%%%%%%%%%%%%%%%%%%%%%%%%%%%%%%%%%
%%%%%%%%%%%%%%%%%%%%
```

```
function dy = ode_six(t,y) % Function which returns the derivatives
% Assumed weights are the following
w_ET=1; w_AT=1;
w_TG=1; w_DA=1; w_TA=1; w_AV=1;
dy=zeros(6,1); % Array initialization to all zeros
% Equations from 66-71
dy(1)=0;
dy(2)=w_ET*y(1)-w_AT*y(5)*y(2);
dy(3)=w_TG*y(2);
dy(4)=0;
dy(5)=w_DA*y(4)-w_TA*y(2)*y(5);
dy(6)=w_AV*y(5);
end
```

**TOR-FLZ8-SnRK1 signaling in favourable conditions:**

1. **With FLZ8**

To predict the role of FLZ8 module in growth and biomass accumulation in favourable condition, we assumed an equal initial value (1 unit each) for TOR and SnRK1 as base. Using the network model described above; we predicted the growth with and without FLZ8 under energy sufficiency (1 unit). The outcome was presented in Fig. 7a.

The equations 35-40 are in the first order form. The array of six unknowns (E, T, S, F, G, B) were assumed as y=[y(1) y(2) y(3) y(4) y(5) y(6)]. Note that the order of the unknowns does not matter. An array of the six derivatives was assumed accordingly, as dy. Consequently, the system of equations 35-40 were rewritten as:

$$dy(1) = 0 \quad (66)$$
$$dy(2) = w_{ET}y(1) - w_{ST}y(3)y(2) \quad (67)$$
$$dy(3) = -w_{ES}y(1)y(3) - w_{TS}y(3)y(2) + w_{FS}y(4) \quad (68)$$
$$dy(4) = w_{TF}y(2) + w_{EF}y(1) - w_{SF}y(3)y(4) \quad (69)$$
$$dy(5) = w_{TG}y(2) - w_{SG}y(3)y(5) \quad (70)$$
$$dy(6) = w_{GB}y(5) \quad (71)$$

A simple MATLAB code with all the descriptions is given below. Note that the symbol "%" should be used in the code for user comments. Some of the comments were omitted for compactness.

```
%% MATLAB code for the solution for favourable conditions with FLZ8
TimeSpan=[0 10]; % Initial time is 0 and the final time is 10
InitialValue=[1 1 1 1 1 1]; % Prescribed initial values for E, T, S, % F, G, B based on equations
66-71
%% Main function
 [t,y]=ode45(@ode_six,TimeSpan,InitialValue); % Ode45 solver

%% Sample plot for the solution
% Plot for B (Biomass), which is the 6th unknown
```

```
plot(t,y(:,6)); % Plotting the 6th column of the matrix [y] in Y-axis
% with time (t) on the X-axis.

%%%%%%%%%%%%%%%%%%%%%%%%%%%%%%%%%%%%%%%%%%%%%%%%
%%%%%%%%%%%%%%%%%%%%%%
% Write here your code to plot any other solution


%%%%%%%%%%%%%%%%%%%%%%%%%%%%%%%%%%%%%%%%%%%%%%%%
%%%%%%%%%%%%%%%%%%%%%%

function dy = ode_six(t,y) % Function which returns the derivatives
% Assumed weights are the following
w_ET=1; w_ST=1; w_ES=1; w_TS=1;
w_TF=0.25; w_EF=0.25; w_SF=0.25; w_FS=0.25;
w_TG=1; w_SG=1; w_GB=1;
dy=zeros(6,1); % Array initialization to all zeros
% Equations from 38-43
dy(1)=0;
dy(2)=w_ET*y(1)-w_ST*y(3)*y(2);
dy(3)=-w_ES*y(1)*y(3)-w_TS*y(3)*y(2)+w_FS*y(4);
dy(4)=w_TF*y(2)+w_EF*y(1)-w_SF*y(3)*y(4);
dy(5)= w_TG*y(2)-w_SG*y(3)*y(5);
dy(6)=w_GB*y(5);
end
```

**2. Without FLZ8**

In this case, the number of unknowns in the equations 14-19 were reduced to five as the variable F was not considered. The system consists of all the first order equations. The variables (E, T, S, G, B) were assumed as y=[y(1) y(2) y(3) y(4) y(5)]. The pathways consisting of FLZ8 were deleted, which caused the corresponding weights to be set to zero. dy reduced to an array of five derivatives. Consequently, the system of equations became:

$$dy(1) = 0 \quad (44)$$

$$dy(2) = w_{ET}y(1) - w_{ST}y(3)y(2) \quad (45)$$

$$dy(3) = -w_{ES}y(1)y(3) - w_{TS}y(3)y(2) \quad (46)$$

$$dy(4) = w_{TG}y(2) - w_{SG}y(3)y(4) \quad (47)$$

$$dy(5) = w_{GB}y(4) \quad (48)$$

The MATLAB code is given below. The same initial conditions, time span and weights as in the previous section were used for comparison. Some of the comments were omitted for compactness.

```
%% MATLAB code for the solution for favourable conditions
%% without FLZ8
TimeSpan=[0 10];
InitialValue=[1 1 1 1 1]; % Initial values for E, T, S, G, B
% based on equations 20-25 deleting the initial value for F
%% Main function
[t,y]=ode45(@ode_five,TimeSpan,InitialValue);
```

```
%% Sample plot for the solution
% Plot for B (Biomass), which is the 5th unknown
plot(t,y(:,5));
%%%%%%%%%%%%%%%%%%%%%%%%%%%%%%%%%%%%%%%%%%%%%%%%%%%%%%%%%%%%%%%%%%%%%%%%%%%%%%%%%%%%%%%%%%%%%%%%%%%
%%%%%%%%%%%%%%%%%%%%%%%%%%%%%%%%%%%%%%%%
% Write here your code to plot any other solution

%%%%%%%%%%%%%%%%%%%%%%%%%%%%%%%%%%%%%%%%%%%%%%%%%%%%%%%%%%%%%%%%%%%%%%%%%%%%%%%%%%%%%%%%%%%%%%%%%%%
%%%%%%%%%%%%%%%%%%%%%%%%%%%%%%%%%%%%%%%%
function dy = ode_five(t,y) % Function which returns the derivatives
% Assumed weights are the following
w_ET=1; w_ST=1; w_ES=1; w_TS=1;
w_TG=1; w_SG=1; w_GB=1;
dy=zeros(5,1); % Array initialization to all zeros
% Equations 44-48
dy(1)=0;
dy(2)=w_ET*y(1)-w_ST*y(3)*y(2);
dy(3)=-w_ES*y(1)*y(3)-w_TS*y(3)*y(2);
dy(4)= w_TG*y(2)-w_SG*y(3)*y(4);
dy(5)=w_GB*y(4);
end
```

**TOR-FLZ8-SnRK1 signaling in unfavourable conditions:**

1. **With FLZ8**

To predict the role of FLZ8 module in stress response, we assumed that the plants are growing in a favourable condition (TOR: 10 units, SnRK1: 1 unit). Using the network model described above; we predicted the stress response and survival in response to sudden stress (1 unit) in models with or without FLZ8. The outcome was presented in Fig. 7b-d.

A similar approach as in the previous sections was used to solve the system of ODE's as IVP. The array of six unknowns (D, T, S, F, R, V) were assumed as u=[u(1) u(2) u(3) u(4) u(5) u(6)]. An array of the six derivatives was assumed as du accordingly. Consequently, the system of equations (26-31) was rewritten as:

$$du(1) = 0 \quad (49)$$
$$du(2) = -w_{DT}u(1)u(2) - w_{ST}u(3)u(2) \quad (50)$$
$$du(3) = -w_{DS}u(1) - w_{TS}u(3)u(2) + w_{FS}u(4) \quad (51)$$
$$du(4) = w_{TF}u(2) - w_{SF}u(3)u(4) \quad (52)$$
$$du(5) = w_{SR}u(3) - w_{TR}u(2)u(5) \quad (53)$$
$$du(6) = w_{RV}u(5) \quad (54)$$

The MATLAB code is given below. Some comments were omitted for compactness.

```
%% MATLAB code for the unfavourable conditions with FLZ8
TimeSpan=[0 10];
InitialValue=[1 10 1 1 1 1]; % Initial values for D, T, S, F, R, V
% based on euations 32-37.
%% Main function
```

```
[t,u]=ode45(@ode_six,TimeSpan,InitialValue);
%% Sample plot for the solution
% Plot for V (Survival), which is the 6th unknown
plot(t,u(:,6));
%%%%%%%%%%%%%%%%%%%%%%%%%%%%%%%%%%%%%%%%%%%%%%%
%%%%%%%%%%%%%%%%%%%%%
% Write here your code to plot any other solution

%%%%%%%%%%%%%%%%%%%%%%%%%%%%%%%%%%%%%%%%%%%%%%%
%%%%%%%%%%%%%%%%%%%%%
function du = ode_six(t,u)
% Assumed weights are the following
w_DT=1; w_ST=1; w_DS=1; w_TS=1;
w_TF=0.25;w_SF=0.25; w_FS=0.25;
w_TR=1; w_SR=1; w_RV=1;
du=zeros(6,1); % Array initialization to all zeros
% Equations 49-54
du(1)=0;
du(2)=-w_DT*u(1)*u(2)-w_ST*u(3)*u(2);
du(3)=w_DS*u(1)-w_TS*u(2)*u(3)+w_FS*u(4);
du(4)=w_TF*u(2)-w_SF*u(3)*u(4);
du(5)= w_SR*u(3)-w_TR*u(2)*u(5);
du(6)=w_RV*u(5);
end
```

**2. Without FLZ8**

The derivatives and the initial values were determined as explained in the previous sections. The MATLAB code is given below:

```
%% MATLAB code for the unfavourable conditions without FLZ8
TimeSpan=[0 10];
InitialValue=[1 10 1 1 1]; % Initial values for D, T, S, R, V
%% Main function
[t,u]=ode45(@ode_five,TimeSpan,InitialValue);
%% Sample plot for the solution
% Plot for V (Survival), which is the 5th unknown
plot(t,u(:,5));
%%%%%%%%%%%%%%%%%%%%%%%%%%%%%%%%%%%%%%%%%%%%%%%
%%%%%%%%%%%%%%%%%%%%%
% Write here your code to plot any other solution

%%%%%%%%%%%%%%%%%%%%%%%%%%%%%%%%%%%%%%%%%%%%%%%
%%%%%%%%%%%%%%%%%%%%%
function du = ode_five(t,u)
% Assumed weights are the following
w_DT=1; w_ST=1; w_DS=1; w_TS=1;
w_TR=1; w_SR=1; w_RV=1;
du=zeros(5,1); % Array initialization to all zeros
du(1)=0;
du(2)=-w_DT*u(1)*u(2)-w_ST*u(3)*u(2);
du(3)=w_DS*u(1)-w_TS*u(2)*u(3);
```

```
du(4)= w_SR*u(3)-w_TR*u(2)*u(4);
du(5)=w_RV*u(4);
end
```

**Supplementary references**


1. Wang, P. *et al.* Reciprocal Regulation of the TOR Kinase and ABA Receptor Balances Plant Growth and Stress Response. *Mol. Cell* **69**, 100-112.e6 (2018).

2. Hindupur, S. K., González, A. & Hall, M. N. The opposing actions of target of rapamycin and AMP-activated protein kinase in cell growth control. *Cold Spring Harb. Perspect. Biol.* **7**, a019141 (2015).

3. Broeckx, T., Hulsmans, S. & Rolland, F. The plant energy sensor: evolutionary conservation and divergence of SnRK1 structure, regulation, and function. *J. Exp. Bot.* **67**, 6215–6252 (2016).

4. Baena-González, E., Rolland, F., Thevelein, J. M. & Sheen, J. A central integrator of transcription networks in plant stress and energy signalling. *Nature* **448**, 938–942 (2007).

5. Xiong, Y. *et al.* Glucose–TOR signalling reprograms the transcriptome and activates meristems. *Nature* **496**, 181–186 (2013).

6. Deprost, D. *et al.* The Arabidopsis TOR kinase links plant growth, yield, stress resistance and mRNA translation. *EMBO Rep.* **8**, 864–870 (2007).

7. Gwinn, D. M. *et al.* AMPK Phosphorylation of Raptor Mediates a Metabolic Checkpoint. *Mol. Cell* **30**, 214–226 (2008).

8. Nukarinen, E. *et al.* Quantitative phosphoproteomics reveals the role of the AMPK plant ortholog SnRK1 as a metabolic master regulator under energy deprivation. *Sci. Rep.* **6**, 31697 (2016).

9. Ling, N. X. Y. *et al.* mTORC1 directly inhibits AMPK to promote cell proliferation under nutrient stress. *Nat. Metab.* (2020). doi:10.1038/s42255-019-0157-1

10. Margalha, L., Confraria, A. & Baena-González, E. SnRK1 and TOR: modulating growth–defense trade-offs in plant stress responses. *J. Exp. Bot.* **70**, 2261–2274 (2019).

11. De Vleesschauwer, D. *et al.* Target of rapamycin signaling orchestrates growth-defense trade-offs in plants. *New Phytol.* **217**, 305–319 (2018).

12. Rodrigues, A. *et al.* ABI1 and PP2CA Phosphatases Are Negative Regulators of Snf1-Related Protein Kinase1 Signaling in Arabidopsis. *Plant Cell* **25**, 3871–3884 (2013).

13. Filipe, O., De Vleesschauwer, D., Haeck, A., Demeestere, K. & Höfte, M. The energy sensor OsSnRK1a confers broad-spectrum disease resistance in rice. *Sci. Rep.* **8**, 3864 (2018).
