## Supplementary Table S2 for "A negative feedback loop of the TOR signaling moderates growth and enables rapid sensing of stress signals in plants"

| **Table S2: Primers used in this study** | |
|---|---|
| **Cloning** | |
| FLZ8 F | ATGCTAAAGAAGAGATCAAG |
| FLZ8 R | TTTAGTATCATTTTCCTCTG |
| pFLZ8 F | CCATGGCTTACTTAATAACAATTATAAAC |
| pFLZ8 R | CTGCAGAATAATAATACTTATATGAATCAC |
| SnRK1α1 F | ATGGATGGATCAGGCACAG |
| SnRK1α1 R | TCAGATCACACGAAGCTCTG |
| SnRK1α1_CD F | ATGGATGGATCAGGCACAGGCAG |
| SnRK1α1_CD R | TCAGAACCAAGGGTGTTGCCGG |
| SnRK1α1_RD F | ATGCAAGCTCATCTTCCGAGGTATT |
| SnRK1α1_RD R | TCAGAGGACTCGGAGCTGAGCAAG |
| SnRK1α1_UBA F | ATGAAGATTGACGAGGAGATTCTC |
| SnRK1α1_UBA R | TCAGTCCAGTATCAGATAGTACGTCAC |
| SnRK1α1_αCTD F | ATGGCAGCTGTTAAGTCGCC |
| SnRK1α1_ αCTD R | TCAGAGGACTCGGAGCTGAGC |
| RAPTOR1B F | ATGGCATTAGGAGACTTAAT |
| RAPTOR1B R | TCATCTTGCTTGCGAGT |
| RAPTOR1B_RNC F | ATGGCATTAGGAGACTTAATGGT |
| RAPTOR1B_RNC R | TCACATAGGGTGAGATATTGGATTAC |
| RAPTOR1B_HEAT F | ATGCTGCCTCCTACGCATCAAC |
| RAPTOR1B_HEAT R | TCAAAGTGCGAATTTTTCTTTCTCTTC |
| RAPTOR1B_WD40 F | ATGGAGCATATTGCAAAATGCC |
| RAPTOR1B_WD40 R | TCATCTTGCTTGCGAGTTGTCG |
| **ChIP qRT-PCR** | |
| pFLZ8_R1 F | CAGTTACTTTTCTCTTTTGTCACACTTTCATA |
| pFLZ8_R1 R | CACTGCTCAATCACACACACACAC |
| pFLZ8_R2 F | GTATTATTATTTGATAGAGAAGCTAAGTGTGTGTC |
| pFLZ8_R2 R | GGAGAAAAGAAGAAGAAACAAACAAAAGAA |
| pFLZ8_R3 F | CTGTTTTGAAAACGTTTTAGTACACTATAATGTG |
| pFLZ8_R3 R | GAGAAAAGGAACTTTAGAAATTTAATGAGAAAG |
| pFLZ8_R4 F | CTCGTATTCATCATCTCTATTTGTCTCCC |
| pFLZ8_R4 R | GAGATCCTTTGCTAGTAAAATAATGTCACA |
| pMCM5_R1 F | AGAAAGAAAGACCCAATAACCAAC |
| pMCM5_R1 R | TCTAAACGAAGAGAGAGAGTGGG |
| pACT7_R1 F | CGTTTCGCTTTCCTTAGTGTTAGCT |
| pACT7_R1 R | CGTTTCGCTTTCCTTAGTGTTAGCT |
| **qRT-PCR** | |
| FLZ8_RT1 F | CCAGAATCATCTCCGGCTATTTC |
| FLZ8_RT1 R | CGCATGTGTAATCCTCCGATAAC |
| FLZ8_RT2 F | ACGAGGCCCAAGACTCTTCTT |
| FLZ8_RT2 R | TCCCCACCCGACAGTTTTC |
| MCM3_RT F | TTCGCCACAAGCGAGATTT |
| MCM3_RT R | CAAAGCCTTGATTTCCTCCATATAC |
| ETG1_RT F | CGGTCTACTTGGGAATGATCACA |

| | |
|---|---|
| ETG1_RT R | CACCTTTGACAGGAGATGCAATAA |
| DIN6_RT F | AACTTGTCGCCAGATCAAGG |
| DIN6_RT R | GGAACACGTGCCTCTAGTCC |
| DIN10_RT F | CGTTGGGATTAAGAACATTGTCA |
| DIN10_RT R | TGCCACACGTAAACATATTTCA |
| TOR_RT F | TTTGATTGGCTTCGCGTAGA |
| TOR_RT R | AACGGCGGCGAAACG |
| RAPTOR1A_RT F | CTCTCCGCCGCCACAA |
| RAPTOR1A _RT R | TGTCTAAGGTCACAGAGCACAAGAGT |
| RAPTOR1B_RT F | TGGTCTCCAATCACCGTTACG |
| RAPTOR1B _RT R | CCGGGAATCACCGTCATC |
| LST8-1_RT F | CGGGATAAAATCGATCGAAAAT |
| LST8-1_RT R | TGTGATCATAGCTAGCCGTAGCAA |
| S6K1_RT F | TCTCTCTCCAGGCTGGTGAAC |
| S6K1_RT R | CGGGAAAAATCGATACAGAATCA |
| S6K2_RT F | GGTAGCAGAGCCTCATCCATCT |
| S6K2_RT R | CAATCGCCAAAAACAATCACAGA |
| SnRK1α1_RT F | GCGCAGATGGTATGCTCAGTAA |
| SnRK1α1_RT R | TGCTGGACTCGTCTCCAAAGT |
| SnRK1α2_RT F | AGGTGAAAATAGCAGAGCATGTTG |
| SnRK1α2_RT R | TTACGACGATTAAGGATTTTGATAGC |
| UBQ10_RT F | TGAAGACTCTCACCGGAAAGACTAT |
| UBQ10_RT R | AGAAGTTCGACTTGTCATTAGAAAGAAA |
| RD29A_RTF | GTGGGCTTTGGTGACGAGTC |
| RD29A_RTR | GTGTCCATTCCAGTTTCAGTCTTC |
| RD29B_RTF | GCGGCGGGCAAAGC |
| RD29B_RTR | AAGCAGTAACAGATCTCGGAGTTTC |
| **Genotyping** | |
| SALK LBb1 | GCGTGGACCGCTTGCTGCAACT |
| SALK RBb1 | TCAGTGACAACGTCGAGCAC |
| SALK_129608_LP | CATGCATCTATAGCTCGGTGAC |
| SALK_129608_RP | GAACCCAAAAGAATCGGACTC |
| SALK_137194_LP | CATGCATCTATAGCTCGGTGAC |
| SALK_137194_RP | GAACCCAAAAGAATCGGACTC |
